# Paint-Lac: A photoactivatable integrator for l-lactate

**DOI:** 10.64898/2026.11.19.767912

**Authors:** Junran Peng, Kei Takahashi-Yamashiro, Takuya Terai, Robert E. Campbell

## Abstract

**SIGNIFICANCE:** This work reports the engineering and characterization of Paint-Lac1, a ligand-modulated, photoactivatable fluorescent integrator for l-lactate, created by combining an mEos-derived photoconvertible protein with an l-lactate binding domain. Upon UV illumination, Paint-Lac1 undergoes l-lactate-inhibited dim-to-bright fluorescence photoactivation to form an orange fluorescent state. Notably, neither the non-photoconverted nor the photoconverted state exhibits fluorescence changes in response to l-lactate concentration, such that the accumulated signal reflects the integrated concentration during the illumination period rather than the concentration at any particular time point. In contrast, the mEos-based Ca^2+^ integrator CaMPARI remains analyte-responsive in both its non-photoconverted and photoconverted states. Analysis of the unique dim-to-bright photoactivation mechanism of Paint-Lac1 reveals that the mEos-derived chromophore can produce distinct photophysical outcomes depending on the specifics of its microenvironment. Paint-Lac1, along with the ratiometric Paint-Lac1-EGFP variant also described in this work, represents a promising new addition to the toolbox of genetically encoded tools and should serve as a template for the development of the next generation of integrators.

**SUMMARY:** Genetically encoded integrators, which convert transient signals into persistent optical signals, can provide unique insights into cell signaling, but the range of target species remains very limited. This work introduces the first photoactivatable integrator for l-lactate, Paint-Lac1, which operates through a ligand-modulated, irreversible β-elimination reaction. Paint-Lac1 is composed of an LldR lactate binding domain fused to an mEos-derived photoconvertible protein. Upon irradiation with 405-nm light, Paint-Lac1 photoconverts to a stable bright orange state, and this process is inhibited by l-lactate. Mechanistic studies reveal that Paint-Lac1 has the same underlying photochemical mechanism as mEos-derived proteins yet exhibits a distinct photophysical outcome, driven by a chromophore environment that stabilizes the phenol form of the chromophore. A ratiometric variant, Paint-Lac1-EGFP, preserves the l-lactate-modulated integrator function and enables biosensor expression-independent quantification. We demonstrate that Paint-Lac1 can be used in mammalian cells to generate a persistent fluorescence signal that is inversely proportional to l-lactate concentration.

## INTRODUCTION

Fluorescent biosensors are indispensable tools in biological research because they enable the visualization of molecular concentrations and various biochemical activities. Among the many formats that exist, genetically encoded biosensors have become especially powerful and they can be expressed directly in living cells and tissues. Most of the genetically encoded biosensors function as indicators, generating transient signals due to the reversible interactions between sensor and target molecules. This reversibility enables real-time monitoring of dynamics of target molecules with high resolution. A prominent example is the GCaMP series^1–7^ of calcium ion (Ca^2+^) indicators, which have been extensively developed over recent decades. GCaMP indicators are composed of circularly permuted (cp) green fluorescent protein (GFP) inserted into a Ca^2+^-binding domain. Late generation members of the GCaMP series are widely used in neuroscience research for visualization of neural activity.

Compared to indicators like GCaMP, integrators produce permanent and irreversible signals from transient cell signals. For example, Ting and coworkers have reported a transcription-based integrator for intracellular Ca^2+^ called FLiCRE^8^ by combining a protease, a Ca^2+^ binding domain, and a caged transcription factor. When Ca^2+^ is increased, the protease is recruited to and activates the transcription factor, which in turn induces expression of a marker protein such as GFP. Such systems enable molecularly informed characterization and manipulation of activated cellular ensembles in a large field of view, which cannot be addressed by conventional indicators. Some other classes of integrators have also been developed^9, 10^ including chemigenetic systems such as Caprola^11^, but the range of single fluorescent protein (FP)-based integrators remains quite limited, even though their compact genetically encoded architecture, straightforward subcellular localization, and strong biocompatibility should make them particularly attractive for users. To date, the only established single FP-based integrator is the Ca^2+^-modulated photoactivatable ratiometric integrator CaMPARI^12–17^. CaMPARI comprises a circularly permuted photoconvertible fluorescent protein (cpmEos2) fused to a Ca^2+^-binding module (**Figure 1**, right). cpmEos2 was derived from Eos-FP^18^, which undergoes green-to-red photoconversion via an irreversible, light-induced β-elimination of the peptide bond between the chromophore moiety (originally derived from His-Tyr-Gly tripeptide) and the preceding Phe residue, a chemical transformation that extends chromophore conjugation and induces a red spectral shift^19^. In CaMPARI, photoconversion is Ca^2+^-dependent: elevated Ca^2+^ increases the fraction of the sensor converted to the red state, allowing Ca^2+^ concentration to be integrated during the illumination period, as read out using the red-to-green ratio^12^ (**Figure 1**, right). The resulting data provide an irreversible record rather than the transient readout produced by conventional indicators, especially important in large-volume imaging and signal accumulation. However, the green and red signals of CaMPARI are themselves dependent on Ca^2+^ concentration (**Figure 1**, right), hence the quantification of integrated Ca^2+^ signals in living tissue is challenging.

**Figure 1.**
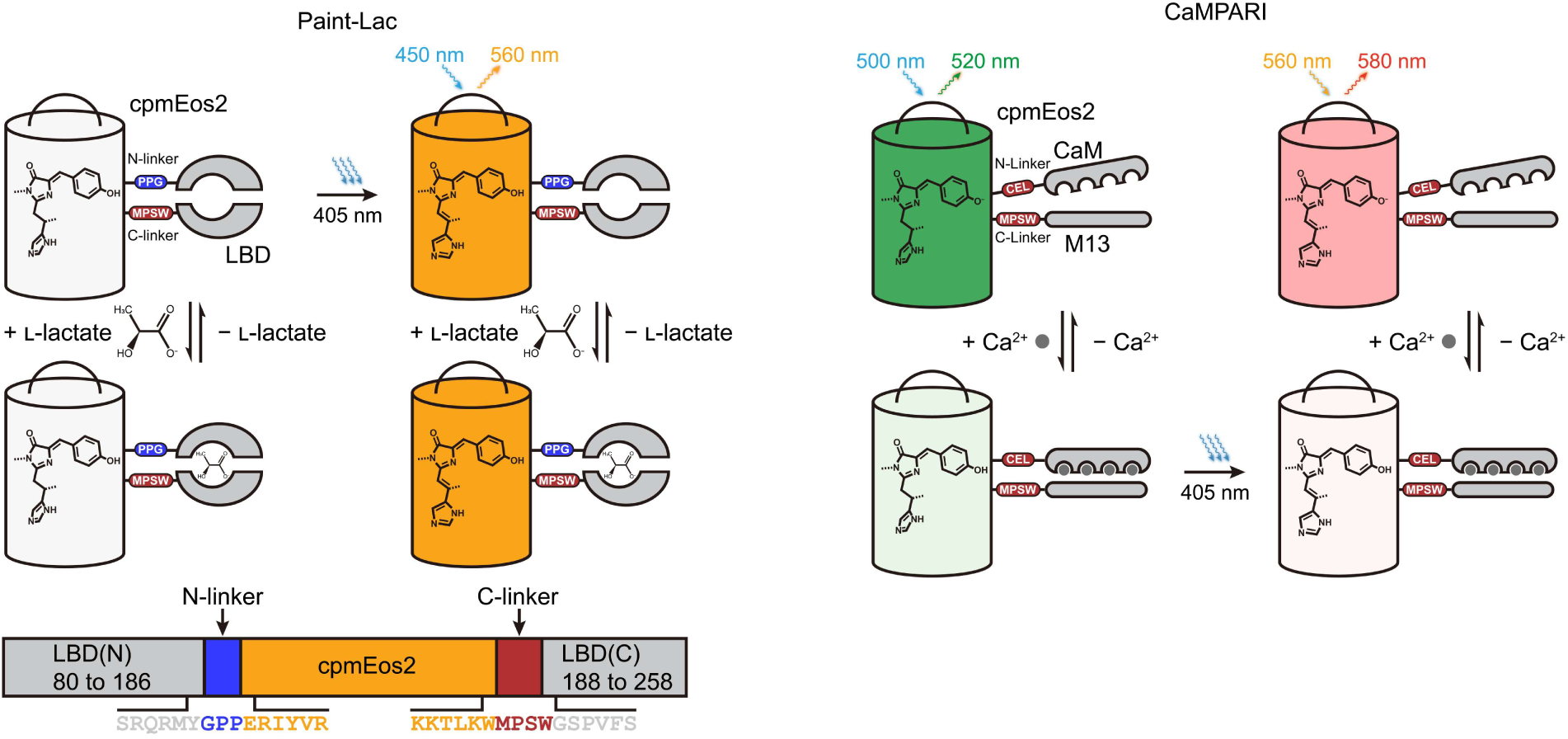
Paint-Lac design and comparison with CaMPARI. Left, schematic representation of Paint-Lac1 structure, target-modulated photoactivation, and responses in dim and bright orange states. The N- and C-terminal parts of LBD (LBD(N), LBD(C)) are shown in gray. Right, schematic representation of CaMPARI structure, target-modulated photoconversion, and responses in green and red states.

The successful development of CaMPARI demonstrated that adapting an irreversible photoconvertible FP within the GCaMP architecture can transform an indicator into an integrator. Building on this concept, we sought to apply a similar strategy to the iLACCO series^20^ of l-lactate indicators. The iLACCO series includes green fluorescent protein (GFP)-based intensiometric biosensors for l-lactate, which were developed by inserting a circularly permutated green fluorescent protein (cpGFP) into the lactate binding domain (LBD) of a transcription factor, LldR^21^. Recent recognition of the diverse physiological roles of l-lactate^22^, including its central role in cellular metabolism and signaling with particular relevance to neural activities^23–26^ and tumor progression^27–29^, has motivated efforts to develop genetically encoded fluorescent biosensors for l-lactate^30–35^, including iLACCO series. Such biosensors enable spatiotemporal analysis of lactate dynamics in living cells and tissues. However, an l-lactate integrator has not yet been reported.

Here, we sought to create an l-lactate integrator by replacing cpGFP in iLACCO1 with cpmEos2 from CaMPARI1 (**Figure 1**). However, we unexpectedly discovered a variant whose cpmEos2 component exhibited unexpected photophysical behavior. Instead of photoconverting from green to red, the variant showed photoactivation from dim to orange in an l-lactate-dependent manner. Taking advantage of this unique finding, we developed Paint-Lac, a photoactivatable integrator for l-lactate that retains CaMPARI’s architecture but displays a distinct photoactivation from a dim state to a bright orange state. This photoactivation was highly l-lactate-dependent, but importantly, fluorescence intensities of bright state was insensitive to l-lactate, providing a basis for simple and robust l-lactate integration (**Figure 1**, left). Paint-Lac1 performs as predicted in *in vitro* characterization with proteins and in mammalian cells, indicating its potential application for recording integrated concentration of l-lactate and for large-volume activity mapping *in vivo*.

## RESULTS

### Development of a photoactivatable integrator for l-lactate, Paint-Lac1

An initial prototype of the l-lactate integrator was constructed by substituting the cpGFP of iLACCO1 with a circularly permuted mEos derived from CaMPARI1 (**Figures 1 and 2A**). Since linker composition is known to play a critical role in FP-based biosensors^36^, we systematically optimized the linkers connecting the two moieties, by only looking at the fluorescence intensity and response to l-lactate (Δ*F*/*F*^min^ = (*F*^l-lactate(−)^ − *F*^l-lactate(+)^) / *F*^l-lactate(+)^) of the initial green state at this early stage of development. We first introduced short flexible linkers (GGS at both the N- and C-termini of cpmEos2) and varied the C-linker length by stepwise deletion, which revealed that a minimal C-linker was optimal (GR-iLACCO0.1d, **Figure S1A,B**). We next tested whether the junction between cpmEos2 and the C-terminal parts of LBD (LBD(C)) could be further shortened by replacing the first two residues (VP) of LBD(C) with a flexible GS sequence and deleting it stepwise. The GS substitution produced the most favorable behavior (GR-iLACCO0.1e, **Figure S1C,D**), and we therefore used GS as the C-linker in subsequent engineering. We then examined the N-linker, where deletion variants showed no improvement over the original GGS sequence.

**Figure 2.**
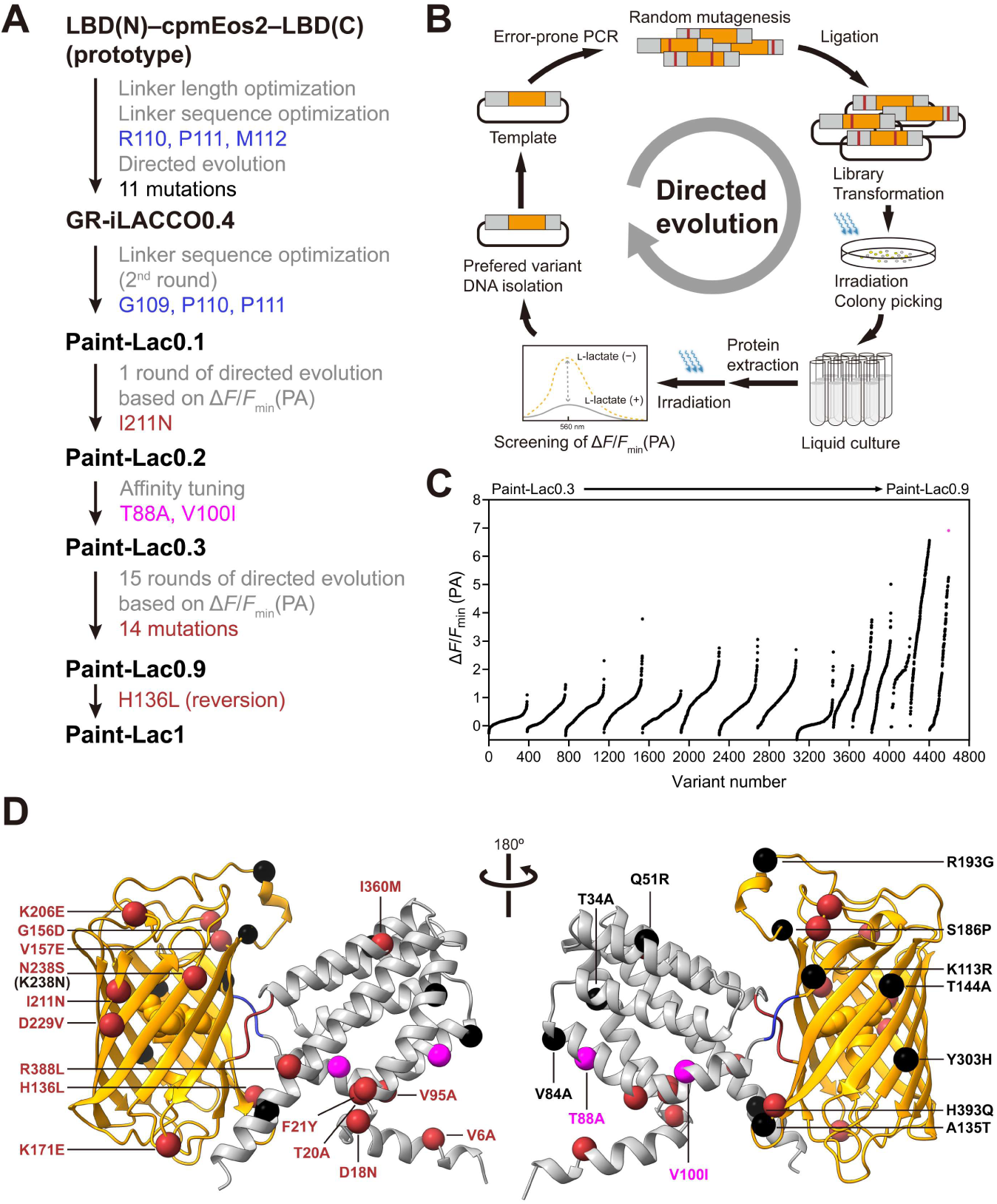
Development of Paint-Lac1. (**A**) Lineage of Paint-Lac variants. Optimization methods are indicated step by step. Mutations introduced in N-linker are shown in blue; mutations introduced by directed evolution to produce GR-iLACCO0.4 are shown in black; mutations introduced by affinity tuning are shown in magenta; mutations introduced by customized directed evolution from Paint-Lac0.3 to Paint-Lac0.9 are shown in red. (**B**) Customized directed evolution protocol for the development of Paint-Lac. After error-prone PCR and building the plasmid library, colonies on agar plates were irradiated with 405 nm light and only the activated colonies were picked for the following culture. The difference in photoactivation with or without l-lactate (Δ*F*/*F*_min_(PA)) was determined using crude protein extract by Bacterial Protein Extraction Reagent (B-PER). Gene encoding the variant with the highest Δ*F*/*F*_min_(PA) was used as template for the next round. (**C**) Δ*F*/*F*_min_(PA) rank plot during the 15 rounds of directed evolution from Paint-Lac0.3 to Paint-Lac0.9. For each round, variants are ranked from the lowest to the highest Δ*F*/*F*_min_(PA) from left to right. (**D**) AlphaFold3^39^ model of Paint-Lac1. Mutations introduced in different stages are labeled with the corresponding colors in (**A**).

These results motivated linker randomization for optimization of the sequence of the N-linker (GGS), yielding a variant with R110, P111, and M112. We then randomized the C-linker (GS), but no improvement was observed. Finally, guided by a reported CaMPARI structure^12^ (**Figure S1E**) showing that the C-terminal residues of cpmEos2 (MPSW) lie close to the chromophore, we randomized these positions as well, although the original sequence again remained among the most favorable. Together, these screens yielded a variant with an inverse response to l-lactate (GR-iLACCO0.3, Δ*F*/*F*^min^ = 0.82 ± 0.048, mean ± s.d., **Figure S1F**), containing an RPM N-linker and a GS C-linker.

To further improve the fluorescence response, we performed 10 rounds of directed evolution on the full-length gene. We expected the sensor to behave similarly to CaMPARI, in which illumination of the protonated chromophore induces green-to-red photoconversion^19, 37^. Under this assumption, a high inverse Δ*F*/*F*^min^ in the green state predicts a strong direct photoconversion response, so we screened libraries produced by error-prone PCR on the basis of green Δ*F*/*F*^min^ (**Figure S2A**). We also irradiated the proteins selected in each round to confirm green-to-red photoconversion, although we did not explicitly screen for l-lactate-dependent photoconversion because the red signal was too dim.

This process yielded a variant containing 11 mutations with a markedly enhanced reversible green-state fluorescence response to l-lactate (GR-iLACCO0.4, Δ*F*/*F*^min^ = 6.4 ± 0.27, **Figure S2B**). However, its photoconversion behavior showed little l-lactate dependence: the red-to-green ratio after photoconversion in the presence or absence of l-lactate differed only slightly (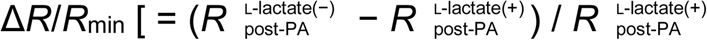 where *R* refers to red-to-green fluorescence ratio] = −0.13 ± 0.060). SDS-PAGE analysis further confirmed that the extent of β-elimination-mediated backbone cleavage was similar with or without l-lactate (**Figure S2C**). Because this behavior is incompatible with the requirements of an integrator, we designated this construct and its predecessors as green-to-red (GR)-iLACCO, reflecting their l-lactate-dependent fluorescence change in both green and red states but their lack of l-lactate-dependent photoconversion. Although red-state Δ*F*/*F*^min^ was not measured for the early variants (GR-iLACCO0.1-0.4), the red states of later variants showed reversible fluorescence responses to l-lactate with similar magnitude to that of their green states (**Figure S3**). This behavior parallels that of GR-GECO^38^, a previously reported highlightable Ca^2+^ indicator based on cpmMaple.

Since the photoconversion of GR-iLACCO0.4 showed no l-lactate dependence, we took a step back and revisited the linker regions. Screening a library in which the three residues of the N-linker were randomized yielded GR-iLACCO0.5, containing L110, P111, and R112. Although GR-iLACCO0.5 still lacked l-lactate-dependent photoconversion, it exhibited a stronger red signal after photoconversion (**Figure S4**), and was therefore retained as the template for the subsequent protein engineering.

Because the primary goal of this work was to engineer a photoactivatable l-lactate integrator, we next examined the N-terminal junction from a different perspective. In iLACCO1, LBD(N) directly connects to cpGFP without an N-linker, and the terminal residue of LBD(N) had been mutated from Leu to Arg during its development^20^. This positively charged Arg might influence the chromophore environment of cpmEos2, therefore, we constructed the second library in which this Arg was removed and the three residues of the resulting N-terminal linker (now residues 109–111) were randomized. Screening this library revealed a distinctive variant that exhibited an unexpected 405-nm-light-induced photoactivation from a dim state to a bright orange state, excitable at 450 nm and emitting at 560 nm (**Figure S5A**). This variant, designated as Paint-Lac0.1a, contains a GPP N-linker. Another variant with a TPP N-linker (Paint-Lac0.1b) showed similar behavior (**Figure S5B**), suggesting that the two Pro residues in the N-linker are likely important for enabling the transition from photoconversion to photoactivation. We note that, using our previously proposed linker definitions^36^, the N-terminal linker length is 1 (that is, the residue aligned with the S142 gatepost residue of mEos2 would be the first P of the GPP linker), and the C-terminal linker length is 4 (that is, the residue aligned with the W139 gatepost is the W immediately before the MPSW linker)

Motivated by the unusual spectral behavior and the possibility of developing a new class of biosensor, we carried out a customized iterative directed evolution starting from Paint-Lac0.1a using a parameter Δ*F*/*F*^min^ (photoactivation, PA) 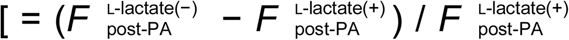, where *F* refers to the fluorescence intensity of 560-nm emission] that reflects the photoactivation bias when irradiated in the presence or absence of l-lactate (**Figure 2B**). The first round introduced a single mutation (I211N) and yielded Paint-Lac0.2 (EC^50^ = 5.37 mM, **Figure S6D**). Before proceeding with further evolution, we incorporated an affinity tuning step to ensure that the biosensor would have an EC^50^ of hundreds of micromolar range which is suitable for cellular and *in vivo* applications. This was achieved by grafting affinity-tuning mutations (T88A, V100I) identified during the development of GR-iLACCO following the strategy established in the iLACCO publication^20^ (**Figure S6A–C**), generating Paint-Lac0.3 (EC^50^ = 101 μM, **Figure S6D**).

Starting from Paint-Lac0.3 we performed 15 rounds of directed evolution that introduced 14 more mutations and produced Paint-Lac0.9 (**Figure 2C,D**). Although Paint-Lac0.9 exhibited a strong photoactivation response, it retained a non-negligible l-lactate dependence in its bright state, which was inconsistent with the permanent signal required for an integrator and contrasted with the behavior of variants isolated after six rounds (**Figure S7**). To eliminate this undesired bright-state response, we constructed libraries that introduced either direct reversions or randomizations at the sites mutated within the cpmEos2 moiety during rounds 6–15. This effort identified a variant carrying an R136L reversion, restoring a Leu residue that had appeared in the third round of evolution and yielding stable bright-state fluorescence while preserving the conserved Δ*F*/*F*^min^(PA). This final construct was designated Paint-Lac1.

### *In vitro* characterization of Paint-Lac1

We evaluated the photophysical and biochemical properties of purified Paint-Lac1 (**Figure 3, Table S1**). Before irradiation, Paint-Lac1 displayed an absorbance peak at 400 nm corresponding to the neutral (protonated) chromophore. After 30 s of 405-nm illumination in the absence of l-lactate, the absorbance peak shifted to 450 nm, whereas the shift was attenuated when l-lactate was present during the illumination (**Figure 3A**). Consistent with these spectral changes, Paint-Lac1 exhibited minimal fluorescence before irradiation and showed an l-lactate-dependent fluorescence enhancement upon photoactivation (Δ*F*/*F*^min^ (PA) = 5.8 ± 0.40, **Figure 3B**). Irradiating Paint-Lac1 across an l-lactate gradient yielded a photoactivation EC^50^ of 16.7 μM (**Figure 3C**). This EC^50^ is lower than the green-state EC^50^ values of GR-iLACCO variants and Paint-Lac0.3 (**Figures S3 and S6**). Although the affinity-tuning mutations were originally grafted with the expectation of producing a similar affinity range, the additional mutations accumulated during directed evolution must have shifted the apparent affinity of the final variant. Both the dim and photoactivated states showed negligible fluorescence changes to l-lactate (**Figure 3D**), demonstrating that Paint-Lac1 produces a permanent signal and functions as a pure integrator, unlike CaMPARI, whose red state remains ligand-responsive after photoconversion.

**Figure 3.**
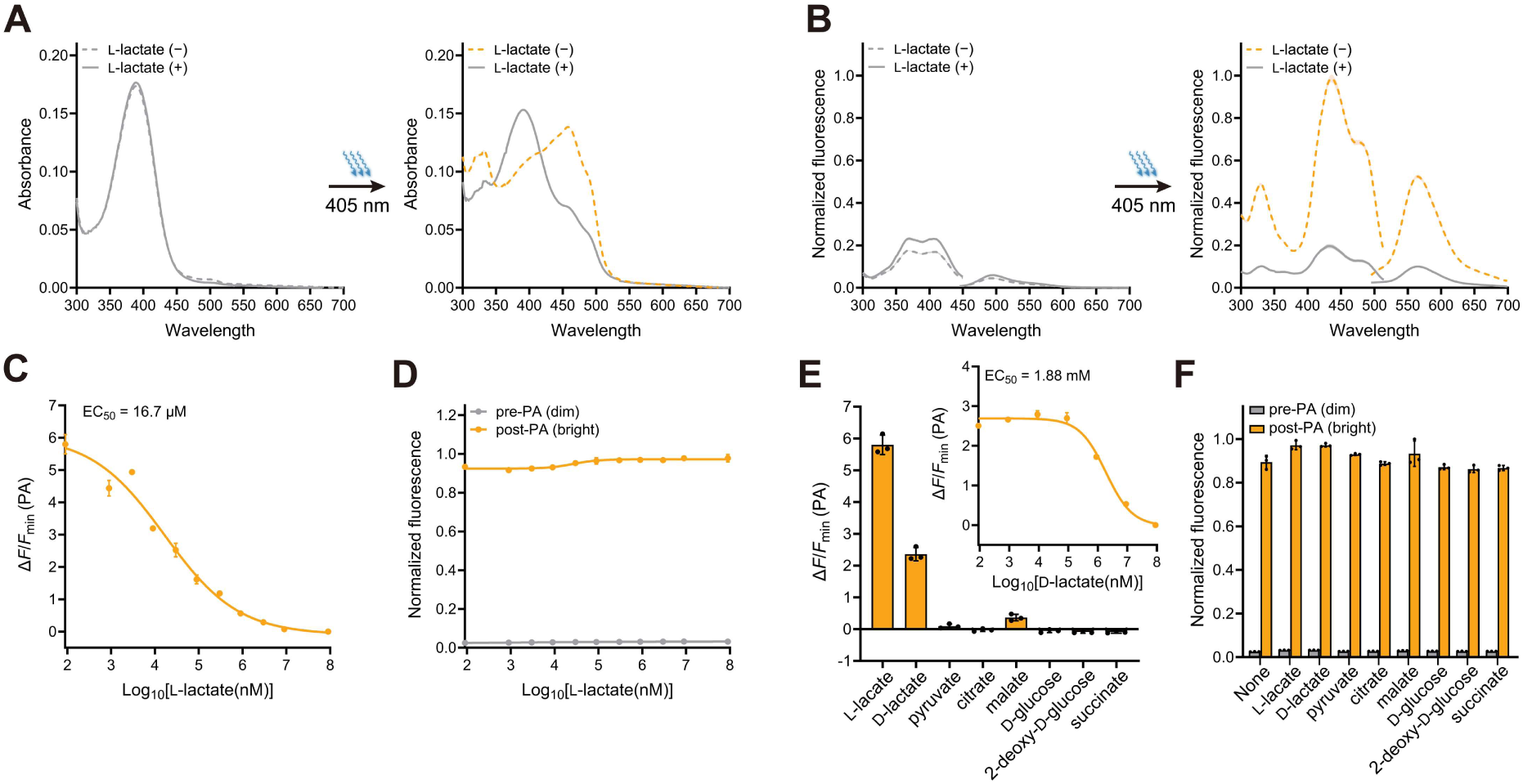
*In vitro* characterization of Paint-Lac1. (**A**) Absorbance spectra of Paint-Lac1 before and after irradiation in the presence (90 mM) and absence of l-lactate. (**B**) Excitation (emission at 500 nm for the dim state and 560 nm for the bright orange state) and emission (excitation at 400 nm for the dim state and 450 nm for the bright orange state) spectra of Paint-Lac1 before and after irradiation in the presence (90 mM) and absence of l-lactate. (**C**) Irradiating Paint-Lac1 across an l-lactate gradient (*n* = 3 technical replicates, mean ± s.d.). (**D**) Dose–response curves of Paint-Lac1 dim and bright orange states for l-lactate (*n* = 3 technical replicates, mean ± s.d.). (**E**) Irradiating Paint-Lac1 across structurally related molecules (90 mM each) and across a ^D^-lactate gradient (*n* = 3 technical replicates, mean ± s.d.). (**F**) Responses toward similar molecules (90 mM each) of Paint-Lac1 dim and bright orange states (*n* = 3 technical replicates, mean ± s.d.).

We next examined the molecular selectivity of Paint-Lac1 using the same photoactivation-based assay described above, irradiating Paint-Lac1 in buffers containing structurally related metabolites. Paint-Lac1 showed strong specificity for l-lactate, with minimal photoactivation responses to the compounds tested (**Figure 3E**). Although Paint-Lac1 responded to D-lactate, its affinity was substantially lower (EC^50^ = 1.88 mM; **Figure 3E** inset), and given that physiological D-lactate concentrations are typically in the tens-of-micromolar range^40^, interference in biological samples is unlikely. We again placed the dim and bright states separately into buffers containing 90 mM of each metabolite (**Figure 3F**). As in the l-lactate gradient experiment, both states exhibited negligible fluorescence changes, confirming that the pure integrator behavior of Paint-Lac1 is maintained even in the presence of high concentrations of structurally similar molecules.

### Investigating the mechanism of photoactivation

The unusual photoactivation behavior of Paint-Lac1 prompted us to investigate its underlying mechanism. Several observations support the conclusion that Paint-Lac1 undergoes the same light-induced β-elimination reaction as CaMPARI, but with some key differences that explain the protein’s unique spectral properties.

First, the chromophore species before and after irradiation appear to be identical to those of CaMPARI. Following the strategy used in the CaMPARI2^13^ study, we digested photoactivated Paint-Lac1 with papain to generate a peptide fragment containing the photoactivated chromophore and analyzed it by mass spectrometry. The mass spectrum of this fragment matched that of the red chromophore from CaMPARI1 (**Figure S8**). Second, SDS-PAGE analysis showed that Paint-Lac1 undergoes light-induced cleavage into two fragments, mirroring the behavior of CaMPARI1, and that the extent of cleavage increased between 30 s and 60 s of 405-nm illumination (**Figure 4A**). Together, these results indicate that Paint-Lac1 employs the same β-elimination chemistry that breaks a peptide bond and extends the chromophore conjugation system in mEos-derived fluorescent proteins.

**Figure 4.**
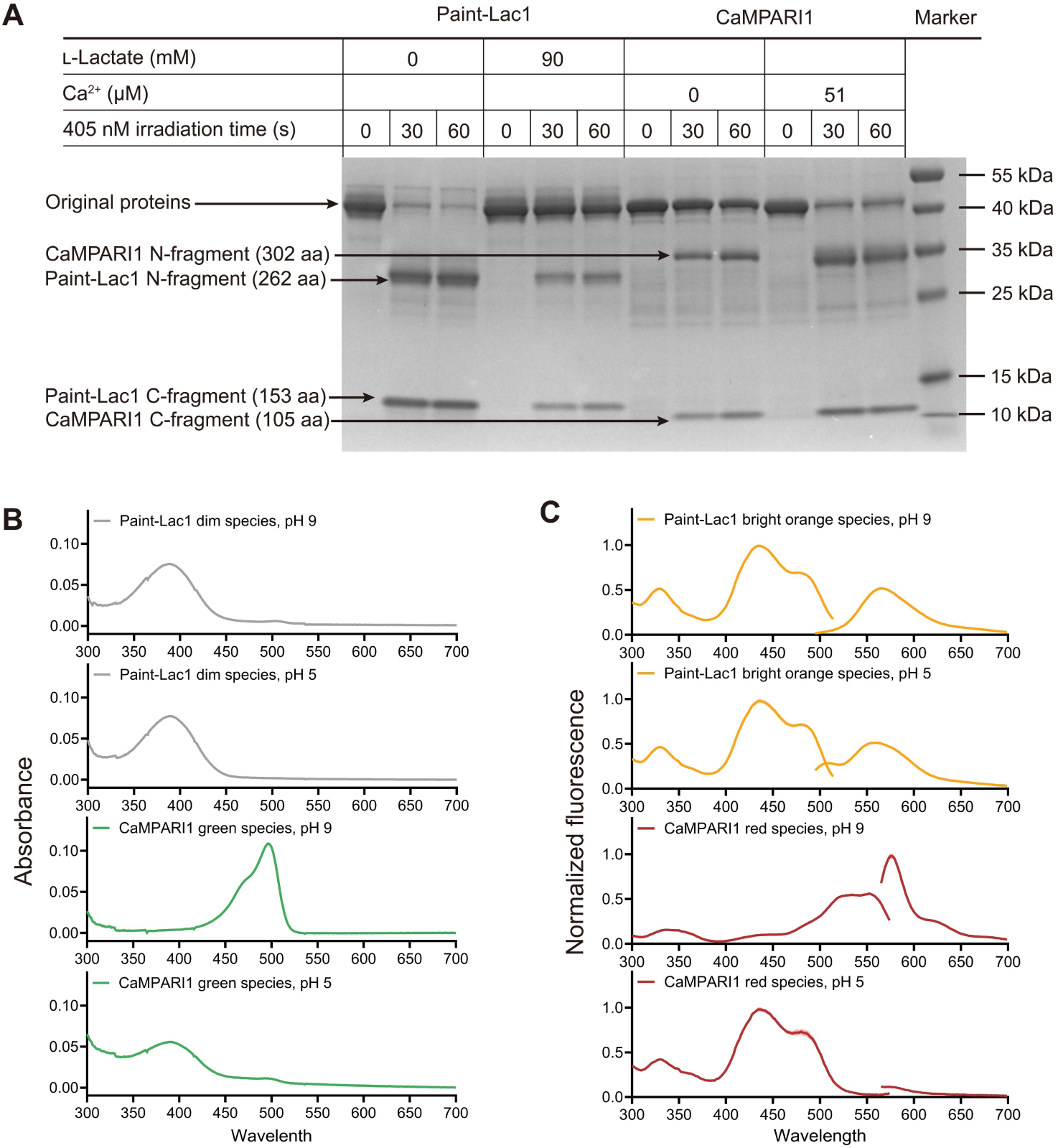
Mechanism investigation of Paint-Lac1. (**A**) SDS-PAGE comparing Paint-Lac1 and CaMPARI1 in terms of their light-induced cleavage. (**B**) Absorbance spectra of Paint-Lac1 dim state and CaMPARI1 green state at pH 9 and 5. (**C**) Excitation (emission at 560 nm for bright orange Paint-Lac1 and 580 nm for red CaMPARI1) and emission (excitation at 450 nm for bright orange Paint-Lac1 and 560 nm for red CaMPARI1) spectra of Paint-Lac1 bright orange state and CaMPARI1 red state at pH 9 and 5.

To understand why this shared reaction produces distinct spectral outcomes, we examined the absorbance and fluorescence of Paint-Lac1 across a range of pH values. From pH 5 to 9, the dim state of Paint-Lac1 consistently exhibited a 400-nm absorbance peak (**Figures 4B and S9A**), matching the neutral (protonated) chromophore observed in CaMPARI1 at pH 5 (**Figures 4B and S9C**). This suggests that the chromophore of the Paint-Lac1 dim species remains predominantly in the neutral form even at high pH, similar to the CaMPARI green state under Ca^2+^-bound conditions, which produces an inverse fluorescence response. Likewise, the bright orange state of Paint-Lac1 remained in the neutral chromophore form across the same pH range (**Figures 4C and S9B**), whereas the red chromophore of CaMPARI1 in the absence of Ca^2+^ shifts between phenol and phenolate states depending on pH (p*K*^a^ = 7.43, **Figures 4C and S9D**). This may be explained by the possibility that the special N-linker GPP, together with the C-terminal MPSW sequence, forms a hydrophobic cage that favors the neutral chromophore (**Figure S10**). Interestingly, CaMPARI1 exhibits a weak 568-nm emission when its red chromophore is excited at 450 nm under acidic conditions (pH 5, **Figure S11**). In contrast, the bright orange state of Paint-Lac1 shows a noticeably stronger 560-nm emission under the same excitation and pH conditions. This difference may arise from a more favorable excited-state proton transfer (ESPT) pathway^41–43^ in Paint-Lac1, enabling the neutral chromophore to emit with a large Stokes shift and producing the characteristic bright orange fluorescence observed after photoactivation.

### Characterization of Paint-Lac1 in mammalian cells

To examine the behavior of Paint-Lac1 in a cellular context, we expressed the biosensor in HeLa cells using a pcDNA3.1^(−)^ plasmid, and evaluated its performance by flow cytometry and fluorescence microscopy. For equilibration between extracellular and intracellular l-lactate, HeLa cells expressing Paint-Lac1 were pretreated with iodoacetic acid to block endogenous production of l-lactate, with nigericin to equalize intracellular and extracellular pH, and with rotenone to inhibit mitochondrial activity. These conditions have been used for characterizing l-lactate indicators^20, 30^. After pretreatment, cells were illuminated with 405-nm light for 30 s either in the absence or presence of 10 mM exogenous l-lactate. Flow cytometry analysis showed that the mean fluorescence intensity (MFI) of cells irradiated without l-lactate was 7.3 ± 0.40-fold higher than that of cells irradiated with l-lactate. (**Figure S12A,B**). This result mirrors the inhibitory effect of l-lactate on photoactivation that we observed with proteins *in vitro*.

To visualize the photoactivation response and the signal stability after irradiation, we performed the fluorescence microscopy on HeLa cells prepared under the same conditions as in the flow cytometry experiments. As shown in **Figures 5A and S13**, Paint-Lac1 fluorescence was initially dim in both representative cells. When the cell 1 was illuminated with 405-nm light, its fluorescence increased markedly, and this activated signal remained unchanged after 5 mM l-lactate was added to the imaging buffer. Conversely, illumination of cell 2 in the presence of 5 mM l-lactate exhibits a noticeably weaker fluorescence increase. On average, the fluorescence of cells activated in the absence of l-lactate was 2.4 ± 1.0 times stronger than in the presence of 5 mM l-lactate (**Figure 5B,C**). The larger fold-change observed in flow cytometry (7.3 ± 0.40-fold) compared with microscopy can be attributed to differences in experimental conditions. Flow cytometry used 10 mM l-lactate, photoactivation of cells in tubes, a broader emission window (570–630 nm) and reported population-averaged MFI, whereas confocal imaging used 5 mM l-lactate, a narrower emission band (540–580 nm), and single-cell illumination and fluorescence readout. Despite these differences, both assays consistently showed that l-lactate attenuates Paint-Lac1 photoactivation. l-lactate also influenced the rate of photoactivation, as fluorescence rose more rapidly when l-lactate was absent, whereas elevated l-lactate slowed the photoactivation (**Figure 5B**). These results clearly demonstrated that Paint-Lac1 can detect l-lactate levels in mammalian cells and record them with stable fluorescent signals after irradiation.

**Figure 5.**
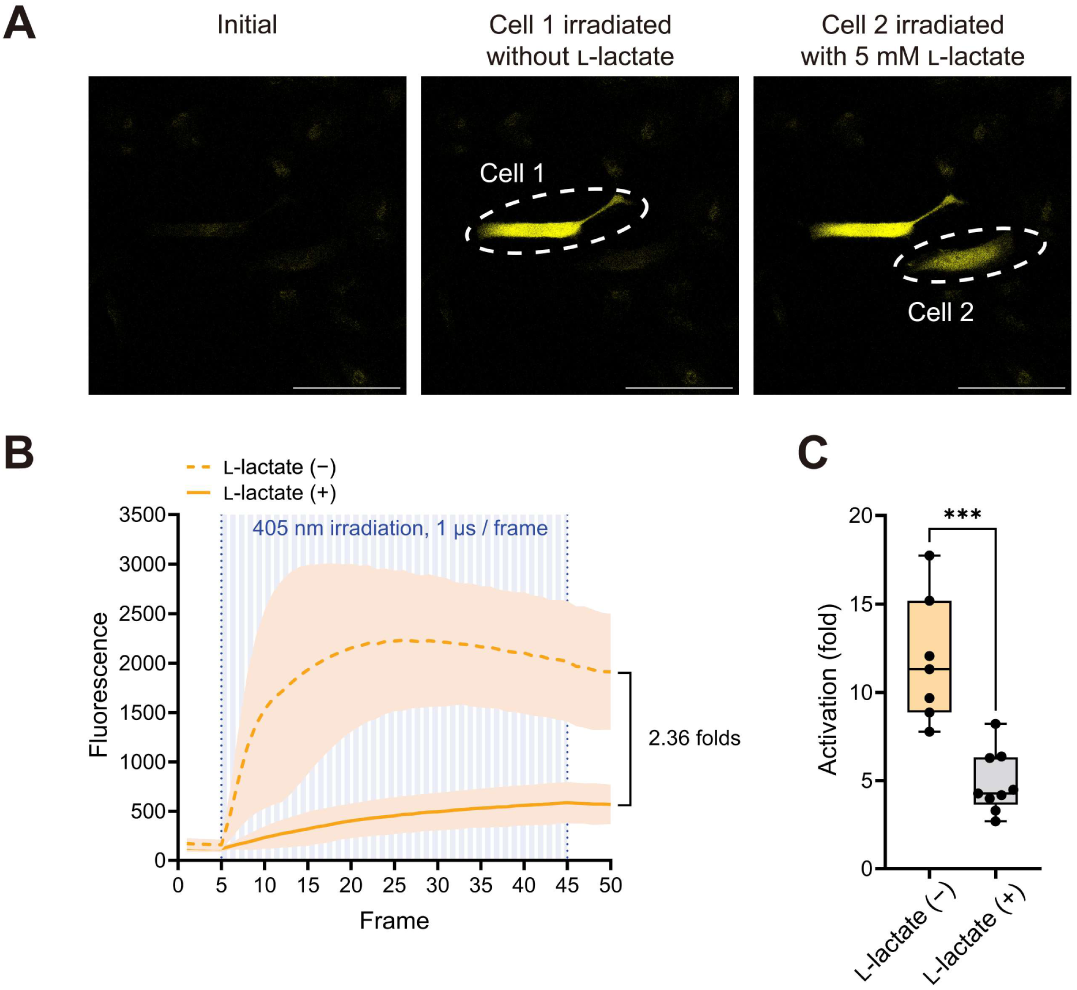
Characterization of Paint-Lac1 in cells. (**A**) Representative images of HeLa cells expressing Paint-Lac1. Cell-selected irradiation was applied with 405-nm light in the absence or presence of 5 mM l-lactate. Scale bars, 100 μm. (**B**) Time courses of the mean ± s.d. of fluorescence of Paint-Lac1 in HeLa cells irradiated without or with 5 mM l-lactate (*n* = 7 and 9 cells, respectively). Irradiation was applied from frame 5 to 45 (1 μs per frame). (**C**) Bar graph showing the activation, defined as *F*_post-PA_ / *F*_pre-PA_ of the cells shown in (**B**) and **Figure S13**. *P* value for technical replicates was calculated using an unpaired nonparametric Mann-Whitney test. \*\*\**P* < 0.05.

### Development and characterization of a ratiometric integrator Paint-Lac1-EGFP

To provide a ratiometric readout that corrects variations in expression and overcomes the very low pre-activation brightness of Paint-Lac1 (as seen in **Figure 5A**, left), we fused an enhanced GFP^44^ (EGFP) as a reference fluorophore to the C-terminus of Paint-Lac1 and Paint-Lac1-EGFP was developed (**Figure 6A**). Because the native EGFP fluorescence was much brighter than the emission from the photoactivated Paint-Lac1 chromophore, the orange peak was initially obscured, preventing reliable ratiometric measurement. Two rounds of directed evolution on the EGFP portion produced a single mutation (D613N) that reduced its brightness sufficiently to allow the two peaks to be clearly distinguished.

**Figure 6.**
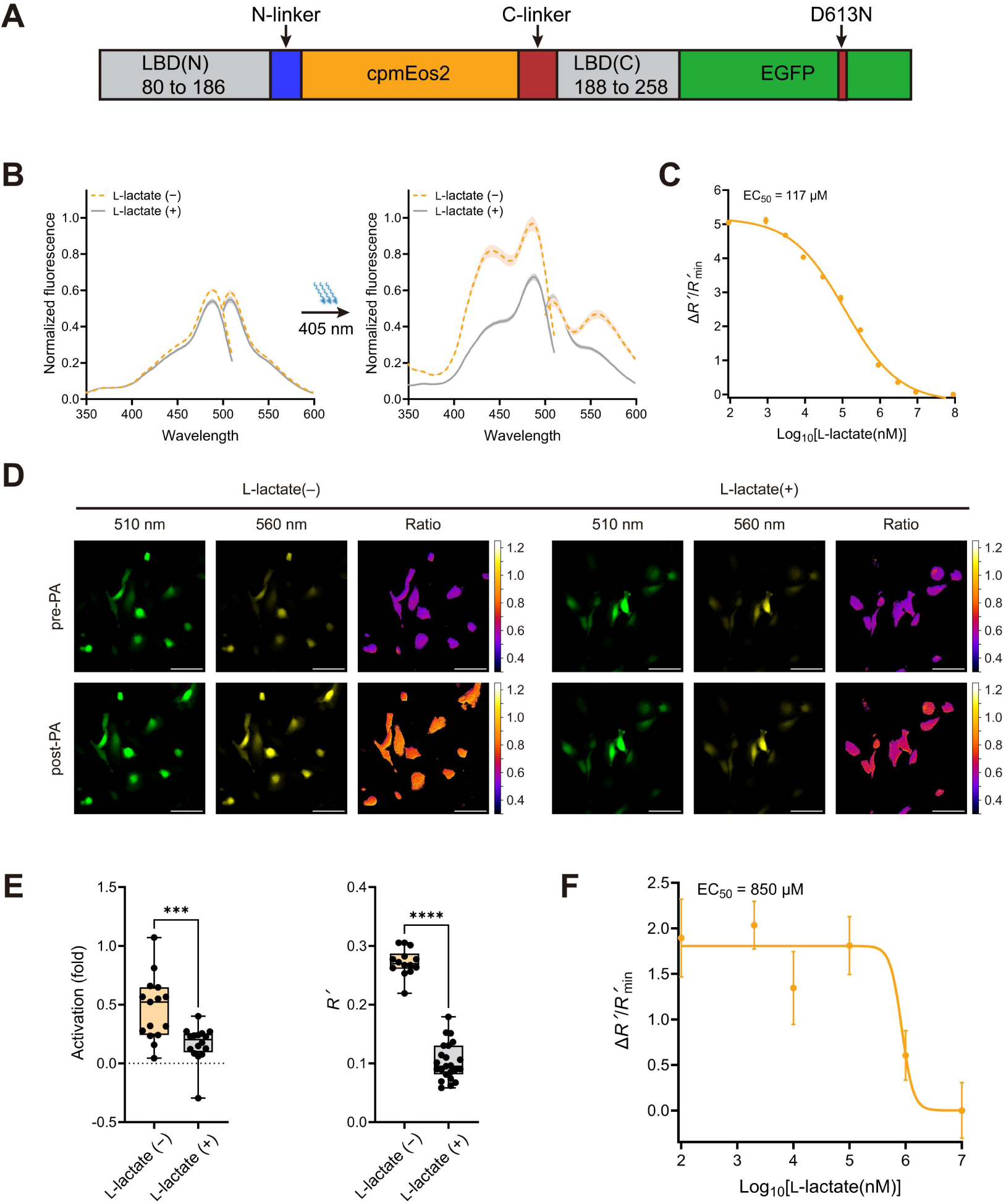
Development and characterization of Paint-Lac1-EGFP. (**A**) Schematic representation of Paint-Lac1-EGFP construct. (**B**) Excitation (emission at 510 nm) and emission (excitation at 450 nm) spectra of Paint-Lac1-EGFP before and after irradiation in the presence (90 mM) or absence of l-lactate. (**C**) Photoactivation of Paint-Lac1-EGFP across an l-lactate gradient. (*n* = 3 technical replicates, mean ± s.d.). (**D**) Representative images of HeLa cells expressing Paint-Lac1-EGFP and irradiated with 405-nm light in the absence or presence of 10 mM l-lactate. Scale bars, 50 μm. (**E**) Left, activation of the orange channel, defined as *F*_post-PA_ / *F*_pre-PA_ for the cells shown in (**D**). Right, ratiometric signal *R*′= *R* – *R*_0_ for the cells shown in (**D**), where *R* = *F*_560_ / *F*_510_ and *R*_0_ is the pre-activation ratio. *P* values for technical replicates were calculated using an unpaired nonparametric Mann-Whitney test (\*\*\**P* < 0.05, \*\*\*\**P* < 0.0001). (**F**) Photoactivation of HeLa cells expressing Paint-Lac1-EGFP across an l-lactate gradient, measured by multichannel fluorescence microscopy (*n* ≥ 12 cells per condition, mean ± s.d.).

*In vitro* measurements showed that only the EGFP portion emitted fluorescence in green before irradiation. After irradiation, the Paint-Lac1 component became photoactivated and the orange peak became visible (**Figure 6B**). Notably, the orange emission displayed the same inverse dependence on l-lactate as observed previously, while the green component was essentially unchanged. Because the EGFP emission tail contributes a non-negligible signal in the 560-nm channel, all ratiometric analyses were performed using a background-subtracted ratio defined as *R*′ = *R* – *R*^0^, where *R* = *F*^560^ / *F*^510^ and *R*^0^ is the pre-activation ratio. Under 450-nm excitation, which provided the most balanced emission pair, the biosensor exhibited a Δ*R*′/*R*′^min^ value of 5.0 ± 0.15. *In vitro* titration with l-lactate revealed an EC^50^ of 117 μM (**Figure 6C**), which is different from that of original Paint-Lac1 (16.7 μM, **Figure 3C**) but falls in the physiological range of l-lactate.

We next expressed Paint-Lac1-EGFP in HeLa cells to evaluate its performance in mammalian cells. The imaging procedure followed the same protocol used for Paint-Lac1, except that fluorescence was collected from both the EGFP channel and the orange channel corresponding to the photoactivated Paint-Lac1 moiety. Representative images illustrating this ratiometric integration of l-lactate in HeLa cells are shown in **Figure 6D**. Comparison of the activation fold in the orange channel with the background-subtracted *R*′ values (**Figure 6E**) indicates that, in contrast to the intensiometric output of Paint-Lac1, Paint-Lac1-EGFP yields a ratio-based readout with smaller standardize deviation due to normalization against cell-dependent expression levels. The measured Δ*R*′/*R*′^min^ value was 1.7 ± 0.22 (EC^50^ = 850 μM, **Figures 6F and S14**). Notably, the apparent affinity differs from the *in vitro* result (**Figure 6C**), which may be due to the increased complexity of the cellular environment. These results show that Paint-Lac1-EGFP preserves the behavior observed *in vitro* and provides a reliable ratiometric signal for quantitative imaging.

## DISCUSSION

This study introduces Paint-Lac1, a genetically encoded integrator for l-lactate, whose underlying mechanism fundamentally differs from the one of CaMPARI. Paint-Lac1 undergoes an irreversible photoactivation and the photoactivation efficiency is inhibited by l-lactate presence in a concentration-dependent manner. Importantly, the bright orange state of Paint-Lac1 is permanently fixed and no longer responds to l-lactate, while both the green and red states of CaMPARI remain Ca^2+^ responsive. As a result, Paint-Lac1 produces a stable, ligand-modulated signal without the interpretive complications inherent to CaMPARI’s reversible indicator function, enabling more reliable integration of transient metabolic events.

Mechanistic characterization showed that Paint-Lac1 undergoes the same light-driven β-elimination reaction that mediates photoconversion in mEos fluorescent proteins. Mass spectrometry of the photoactivated chromophore and SDS-PAGE analysis of irradiated protein both support this conclusion. Despite sharing this underlying chemistry with CaMPARI, Paint-Lac1 displays distinct spectral properties. Measurements across a range of pH values indicate that both the dim and bright chromophore species remain predominantly in the neutral state, in contrast to the phenol–phenolate equilibrium observed in CaMPARI. One possible explanation is that the GPP N-linkers and MPSW C-terminal sequence create a more hydrophobic chromophore environment that stabilizes the neutral form. In CaMPARI, the Ca^2+^ dependence of photoconversion arises from favoring the protonated phenol species, which is more reactive in the β-elimination reaction. In Paint-Lac1, the l-lactate dependence may reflect a broader modulation of the reaction pathway, potentially involving ligand-dependent changes in the chromophore distortion and the proton transfer. The long-Stokes-shift emission of the bright state further suggests that excited-state proton transfer (ESPT) occurs in Paint-Lac1. Together, these observations illustrate how differences in the chromophore environment can reshape the photophysical outcome of a shared chemical reaction.

In mammalian cells, Paint-Lac1 preserved its l-lactate-dependent photoactivation and produced a stable fluorescence signal after illumination. Flow cytometry analysis showed that l-lactate suppresses photoactivation at the population level, and cell imaging with fluorescence microscopy confirmed that the activated orange fluorescence remains stable even when l-lactate levels are altered after illumination. These results demonstrate that Paint-Lac1 can reliably integrate transient l-lactate dynamics into a persistent fluorescence signal in living cells.

To expand the utility of the biosensor in terms of quantification, we developed Paint-Lac1-EGFP, a ratiometric variant that incorporates an internal fluorescence reference. Directed evolution of the EGFP portion produced a balanced pair of emission peaks that enabled robust ratiometric measurements upon 450-nm excitation. Paint-Lac1-EGFP retained the key features of the original biosensor, including the l-lactate-inhibited photoactivation and the permanence of the activated orange state, while substantially improving quantitative performance in both *in vitro* and in mammalian cells.

Together, Paint-Lac1 and its ratiometric variant establish a new mechanism for genetically encoded integrators for metabolites, in which ligand binding modulates an irreversible photoactivation reaction. The mechanistic insights gained here, particularly the influence of the chromophore environment on emission properties, highlight how local structural features can shape the photophysical outcome of a shared chromophore and point to potential strategies for tuning integrator behavior through targeted modifications of the surrounding protein matrix. Looking forward, the stable and ligand-dependent activation of Paint-Lac1 may make it well suited for *in vivo* imaging applications aimed at integrating physiological l-lactate in animal models.

### Limitations of the study

Development of an affinity-tuned series of Paint-Lac1-EGFP variants would further expand the utility of this platform, enabling quantitative imaging across cell types that span a wide range of l-lactate concentrations. Structural and spectroscopic analysis will be important for elucidating how lactate binding inhibits the photoactivation process, thereby enabling more predictive engineering.

## Supporting information

Supplementary information for Paint-Lac

## RESOURCE AVAILABILITY

### Lead contact

Requests for further information and resources should be directed to and will be fulfilled by the lead contact, Robert E. Campbell.

### Materials availability

- Paint-Lac1 and Paint-Lac1-EGFP in pBAD and pcDNA3.1^(−)^ vectors have been deposited to [Addgene, name and catalog number or unique identifier].
- Additional materials and reagents generated in this study are available upon a reasonable request from the lead contact and may require a completed Material Transfer Agreement.

### Data and code availability

- This paper does not report original code.
- Any additional information required to reanalyze the data reported in this paper is available from the lead contact upon request.

## ACKNOWLEDGMENTS

We thank Dr. Yusuke Nasu for helpful discussions and for his earlier contributions to lactate biosensor development. This work was supported by grants from the Japan Society for the Promotion of Science (JSPS) KAKENHI (19H05633, 22H04743, 24H02267, and 24H00489 to R.E.C., 21H00273 and 23H02101 to T.T.), JST CREST (JPMJCR25T3), and the Mitsubishi Foundation. J. P. is supported by JST SPRING (JPMJSP2108) and the World-Leading Innovative Graduate Program for Materials Research, Information, and Technology (MERIT-WINGS).

## AUTHOR CONTRIBUTIONS

J.P. designed and performed all experiments unless otherwise noted. K.T-Y. supervised and assisted with confocal microscopy and flow cytometry experiment, and performed flow cytometry data analysis. K.T-Y., T.T., and R.E.C. supervised the research. J.P. wrote the manuscript with guidance and revisions from K.T-Y., T.T., and R.E.C..

## DECLARATION OF INTERESTS

The authors declare no competing interests.

## DECLARATION OF GENERATIVE AI AND AI-ASSISTED TECHNOLOGIES IN THE WRITING PROCESS

During the preparation of this work, the authors used Microsoft Copilot, ChatGPT5.5 and earlier versions (OpenAI) in order to improve the English language and overall readability. After using this tool, the authors carefully reviewed, revised, and edited the content as needed, and take full responsibility for the final version of the manuscript.

## SUPPLEMENTAL INFORMATION

**Document S1. Figures S1–S16, Tables S1, and supplemental references**

## STAR★METHODS

### KEY RESOURCES TABLE

| REAGENT OR RESOURCE | SOURCE | IDENTIFIER |
|---|---|---|
| Bacterial and virus strains |  |  |
| *E. Coli* DH10B | Thermo Fisher Scientific | Cat #18290015 |
| Chemicals, peptides, and recombinant proteins |  |  |
| Sodium L-lactate | Sigma-Aldrich | Cat #71718 |
| MOPS | Dojindo Laboratories | Cat #078969 |
| Potassium chloride | FUJIFILM | Cat #168-22111 |
| Sodium chloride | FUJIFILM | Cat #190-13921 |
| Calcium chloride dihydrate | FUJIFILM | Cat #038-19735 |
| Manganese(II) chloride solution | Sigma-Aldrich | Cat #M1787 |
| Imidazole | FUJIFILM | Cat #099-00013 |
| Disodium hydrogenphosphate | FUJIFILM | Cat #042-30055 |
| Ethylene glycol bis(β-aminoethylether)-N,N,N′,N′-tetraacetic acid (EGTA) | Nacalai Tesque | Cat #08907-42 |
| Ethylenediaminetetraacetic acid (EDTA) | Nacalai Tesque | Cat #15105-35 |
| Trisodium citrate | FUJIFILM | Cat #203-13605 |
| Sodium tetraborate | FUJIFILM | Cat #192-01455 |
| Sodium D-lactate | Sigma-Aldrich | Cat #71716 |
| Sodium pyruvate | FUJIFILM | Cat #191-03061 |
| DL-Malic acid | FUJIFILM | Cat #135-00562 |
| D-Glucose | FUJIFILM | Cat #046-31175 |
| 2-deoxy-D-Glucose | FUJIFILM | Cat #046-06483 |
| Succinic acid | FUJIFILM | Cat #190-04332 |
| L-Arabinose | Fluorochem | Cat #078969 |
| Iodoacetic acid | FUJIFILM | Cat #091-00492 |
| Nigericin Sodium Salt | FUJIFILM | Cat #149-07261 |
| Rotenone | Tocris Bioscience | Cat #3616 |
| Tris-base | Nacalai Tesque | Cat #35406-91 |
| Glycine | FUJIFILM | Cat #072-05285 |
| Glycerol | Nacalai Tesque | Cat #17018-25 |
| Sodium dodecyl sulfate (SDS) | FUJIFILM | Cat #199-07141 |
| Ampicillin | FUJIFILM | Cat #014-23302 |
| Bromophenol Blue | FUJIFILM | Cat #029-02912 |
| Acetic acid | FUJIFILM | Cat #019-08175 |
| Sodium hydroxide | FUJIFILM | Cat #194-18865 |
| Potassium hydroxide | FUJIFILM | Cat #168-21815 |
| Experimental models: Cell lines |  |  |
| HeLa cells | Japanese Collection of Research Bioresources | JCRB9004 |
| Recombinant DNA |  |  |
| pBAD vector | Invitrogen | Cat #V43001 |
| pcDNA3.1<sup>(−)</sup> vector | Invitrogen | Cat #V79520 |
| pBAD-iLACCO1 | Hario et al., 2024 | N/A |
| pcDNA3-CaMPARI | Fosque et al., 2015 | Addgene #60421 |
| EGFP | Cormack et al., 1996 | N/A |
| pBAD-Paint-Lac1 | This work | Addgene # |
| pBAD-Paint-Lac1-EGFP | This work | Addgene # |
| pcDNA3.1<sup>(−)</sup>-Paint-Lac1 | This work | Addgene # |
| pcDNA3.1<sup>(−)</sup>-Paint-Lac1-EGFP | This work | Addgene # |
| Software and algorithms |  |  |
| cellSens software | Olympus | https://evidentscientific.com/ja/products/software/cellsens |
| NIS-Elements software | Nikon | https://www.microscope.healthcare.nikon.com/ja_JP/products/software/nis-elements |
| ImageJ | National Institutes of Health | https://imagej.net/ij/ |
| GraphPad Prism 10 | Graph Pad Software | https://www.graphpad.com |
| FlowJo v10 | BD | https://www.flowjo.com/ |
| Other |  |  |
| Bacterial Protein Extraction Reagent (B-PER) | Thermo Fisher Scientific | Cat #78248 |
| Ni-NTA agarose | FUJIFILM | Cat #141-09764 |

### EXPERIMENTAL MODEL AND STUDY PARTICIPANT DETAILS

#### Cell lines

HeLa cells were cultured in high-glucose Dulbecco’s modified Eagle’s medium (DMEM, Nacalai Tesque, # 16971-55) supplemented with 10% fetal bovine serum (FBS, Sigma-Aldrich, # F7524) and 100 μg/mL penicillin-streptomycin (Nacalai Tesque, #26253-84), at 37C and 5% CO2 in a humified incubator.

### METHOD DETAILS

#### General materials

pBAD plasmids encoding iLACCO1 and CaMPARI1 were used as the template. Phusion high-fidelity DNA polymerase (Thermo Fisher Scientific, #F530) was used for normal PCR amplification, and Taq DNA polymerase (New England Biolabs, #M0273) was used for error-prone PCR. Restriction endonucleases XhoI and HindIII (Thermo Fisher Scientific, #FD0694 and #FD0504), Gibson Assembly Master Mix (New England Biolabs, #E2611), T4 polynucleotide kinase (PNK) (Thermo Fisher Scientific, #EK0031) and T4 ligase (Thermo Fisher Scientific #EL0014) were used for plasmid construction. Products of PCR and restriction endonucleases digestions were purified using agarose gel electrophoresis followed by extraction by GeneJET Gel Extraction Kit (Thermo Fisher Scientific, #K0692). Plasmid DNA was extracted by GeneJET Plasmid Miniprep Kit (Thermo Fisher Scientific, #K0503) and analyzed by DNA sequence service of FASMAC Co. Ltd. The transformation of electrocompetent *E. coli* DH10B was performed with the plasmids using electroporation (Bio-rad). The fluorescence spectra were recorded by Spark spectrometer (TECAN). The absorbance was measured by UV spectrophotometer (Shimadzu, UV-1800). Quantum yield measurements were performed using a Quantaurus-QY system (Hamamatsu, C9920-02G).

#### Engineering of Paint-Lac

Prototype of Paint-Lac was generated by inserting cpmEos domain from CaMPARI1 into the LldR-LBD of iLACCO1 in a pBAD vector using Gibson assembly. For linker length optimization, cpmEos2 fragments with varying lengths of linkers were amplified by PCR and inserted into LldR-LBD using Gibson assembly. Site-specific mutagenesis was performed by introducing mutated or NNK codons at the 5′ end of the forward primer, with three additional upstream nucleotides to prevent sequence shortening during PCR. Reverse primer was designed so that its 5′ end abutted the 5′ end of the forward primer without any overlap, enabling PCR amplification of a full-length linear blunt-ended product with mutations. PCR products were digested with DpnI (Thermo Fisher Scientific, #FD1704) to remove the template plasmid, purified by gel extraction, phosphorylated with T4 PNK, and circularized with T4 ligase. Directed-evolution libraries were generated by error-prone PCR applying 10 μM MnCl^2^ and imbalanced dNTP (a final concentration of 0.2 mM dATP, 0.2 mM dTTP, 1 mM dGTP, and 1 mM dCTP), introducing one to three mutations per gene, followed by cloning into the pBAD vector by restriction digestion and ligation. For both site-saturation and directed evolution libraries, genes were transformed into *E. coli* DH10B and primary screening was performed on LB agar plates, where approximately 2 × 10^3^ colonies were visually inspected after a 4–6 h maturation period. Fluorescent colonies were selected for GR-iLACCO libraries, whereas Paint-Lac libraries were irradiated at 405 nm for 40 seconds by an LED Flood Controller (LOCTITE) and only photoactivated bright colonies were chosen. A total 192–384 selected colonies were transferred into 96-deep-well plates for expression in 1 mL TB medium, and crude lysates were prepared using B-PER. Lysates were diluted into l-lactate(−) or l-lactate(+) buffer and measured in Nunc MicroWell 96-Well Optical Bottom Plates (Thermo Fisher Scientific). The top 8–10 variants from each round were advanced to confirmational assays. Plasmids from selected variants were purified, sent for sequencing, re-transformed, expressed in 5 mL TB cultures and assayed under identical conditions. Amino acid sequences of the key variants are shown in **Figure S15**.

#### Protein purification

Paint-Lac variants and CaMPARI1 in pBAD vectors containing a N-terminal poly His (6×) tag were expressed in *E. coli* DH10B. A single colony was picked into 10 mL of TB containing 100 μg/mL of ampicillin and cultured overnight at 37 °C by shaking at 250 rpm. The cultures were transferred into 200 to 500 mL of ampicillin-supplemented TB and shaken at 37 °C until the OD^600^ of the culture reached 0.6. L-Arabinose (4 mg/L) was then added to induce protein expression, and the culture was shaken at room temperature for 8 to 12 hours. The next day, cells were harvested by centrifugation at 4,000 rpm for 10 minutes at 4 °C. The resulting pellet was resuspended in 8 mL of binding buffer (20 mM disodium hydrogenphosphate, 500 mM NaCl, and 20 mM imidazole, pH 7.2). The suspension was transferred to a 10 mL beaker and sonicated on ice using a cell homogenizer (BRANSON, Sonifier 250) with an output control of 4 and a duty cycle of 50% for 2 minutes, repeated twice. The lysate was centrifuged at 10,000 rpm for 30 minutes at 4 °C. The supernatant was collected and filtered using a 0.45 μm filter unit (Merck Millipore), then loaded onto a Pierce Centrifuge Column (Thermo Scientific) containing 1.5 mL of Ni-NTA agarose resin. The column was rotated at 4 °C for 20 minutes to allow protein binding, then washed four times with 10 mL of wash buffer (20 mM disodium hydrogenphosphate, 500 mM NaCl, and 20 mM imidazole, pH 7.2). Elution was performed five times using 1 mL of elution buffer (20 mM disodium hydrogenphosphate, 500 mM NaCl, and 250 mM imidazole, pH 7.2). The final four eluted fractions were pooled and concentrated using a 50 mL 30 kDa centrifugal filter unit (Merck Millipore, UFC9030), and the buffer was exchanged with l-lactate(−) buffer (30 mM MOPS, 100 mM KCl, pH 7.2). Protein concentration was determined by NanoPhotometer (Implen, NP80), setting protein molecular weight (MW) and ε^280^ according to the protein parameters.

#### *In vitro* characterization

To prepare the bright orange species of Paint-Lac and red species of CaMPARI, protein samples in l-lactate(−) buffer or Ca^2+^(+) buffer (30 mM MOPS, 100 mM KCl, 10 mM EGTA, 10 mM CaCl^2^, pH 7.2), respectively, were irradiated for 40 seconds by LED Flood Controller at 405 nm (energy: 490 mW/cm^2^). When the protein samples need to be irradiated with specific buffers, 10 μL of the proteins were mixed with 90 μL of the buffers in 96-well optical bottom plates and then irradiated by 405-nm light for 40 seconds. For absorbance and fluorescence excitation/emission spectra, l-lactate(−) buffer and l-lactate(+) buffer (30 mM MOPS, 100 mM KCl, 100 mM l-lactate, pH 7.2) were used. To measure EC^50^, buffers with l-lactate concentration gradient ranging from 0 to 100 mM were prepared by diluting l-lactate(+) buffer using l-lactate(−) buffer. To test specificity, buffers (pH 7.2) containing 100 mM structurally related molecules, 30 mM MOPS, and 100 mM KCl were used. For pH titration, pH buffer (30 mM MOPS, 30 mM trisodium citrate, 30 mM sodium borate, 100 mM KCl, and either no l-lactate or 10 mM l-lactate, with pH ranging from 5 to 10) was used to dilute protein solutions. All measurements were conducted at room temperature.

#### SDS-PAGE

Purified protein samples were treated as described in the Results section. A 20 µL aliquot of each sample was mixed with 20 µL of 2× sample buffer (0.125 M Tris-base, 4% (w/v) SDS, 20% (v/v) glycerol and 0.01% (w/v) bromophenol blue) and boiled at 98 °C for 10 minutes. 12.5% Real Gel Plate (BIO CRAFT, #SDG-538) or 5–12% SuperSep Ace gel (FUJIFILM, #192-15201) was placed in an electrophoresis chamber (BIO CRAFT) and was filled with running buffer (25 mM Tris-base, 192 mM Glycine and 0.1% (w/v) SDS). 20 µL of samples and PageRuler Prestained Protein Ladder (Thermo Fisher Scientific, #26617) were loaded into wells. Electrophoresis was run at 120 V for 1 to 1.5 hours. Gels were stained with Coomassie Brilliant Blue (CBB, Nacalai Tesque, #04543-51) overnight and were washed with water several times.

#### Mass spectra of peptides containing chromophores

Purified Paint-Lac1 and CaMPARI1 proteins were irradiated at 405 nm in l-lactate(−) buffer or Ca^2+^(+) buffer, respectively, for 80 seconds using the LED Flood Controller. The irradiated proteins were concentrated using a 500 µL 10 kDa centrifugal filter unit (Merck Millipore, #UFC5010), mixed with 2× sample buffer (0.125 M Tris-base, 4% (w/v) SDS, 20% (v/v) glycerol and 0.01% (w/v) bromophenol blue), and loaded in a 12% Real Gel Plate (BIO CRAFT). SDS-PAGE was run at 120 V for 1.5 hours. Gel bands with yellow coloration were directedly excised without CCB staining, shredded into 3 mL of elution buffer (50 mM Tris-base, 150 mM NaCl, 0.1 mM EDTA, pH 7.5), and shaken at 37 °C overnight. The supernatant was collected after centrifugation and concentrated using a 500 µL 10 kDa centrifugal filter unit (Merck Millipore, #UFC5010). The protein fragments were digested by papain at 37 °C for 5 h followed by incubation at room temperature for 24 h. Because the N-terminal peptide bound to the HYG-formed chromophore has been cleaved by light-induced β-elimination and the papain protease targets a downstream hydrophobic valine residue, the expected digestion product is the HYG-formed chromophore with two downstream amino-acid residues (NR, **Figure S8**). The orange-yellow solutions were desalted and concentrated by a MonoSpin C18 column (GL Sciences). The products were diluted with MeOH to equal volumes and injected into an electrospray ionization (ESI) mass spectrometer (Bruker, compact system) to obtain mass spectra of the chromophore-containing peptides.

#### Imaging of Paint-Lac1 in HeLa cells

The gene encoding Paint-Lac1 was amplified by PCR, digested with restriction enzymes and ligated into a pcDNA3.1^(−)^ vector. HeLa cells were transfected with pcDNA3.1^(−)^-Paint-Lac1 mixed with polyethylenimine (PEI, Polysciences, #24765) in Opti-MEM I (Gibco, #31985070). Cell imaging was performed within 48 h after transfection. Before imaging, cells were incubated with 500 μM iodoacetic acid at 37 °C for 15 min. After washing with Hank’s balanced salt solution (HBSS, Nacalai Tesque, #09735-75) supplemented with 10 mM HEPES (Nacalai Tesque, #17557-94), cells were maintained in 1.9 mL HBSS containing 10 mM HEPES, 10 μM nigericin, and 2 μM rotenone during imaging. When required, 100 μL HBSS containing 10 mM HEPES and 100 mM l-lactate was added to the imaging dish to reach a final concentration of 5 mM l-lactate. Cell-specific light stimulation and fluorescence imaging were conducted on an AXR-NSPARC confocal based super resolution microscope (Nikon). NIS-Elements software was used for the imaging. A PLAN APO λD 20× OFN25 DIC N2 objective lens was used with a confocal zoom factor of 3×. Cells were excited with a 488-nm laser at 19.6% power with a dwell time of 0.5 μs, and the emission was collected using a GaAsP detector from 540 to 580 nm with a gain 58.9%. Cell-selected irradiation was achieved within a defined region of interest (ROI) using a 405-nm laser at 10% power, scanning for 1 μs per pixel for one cycle in a repeated irradiation-imaging loop. Fluorescence images were analyzed with ImageJ software.

#### Flow cytometry analysis of HeLa cells expressing Paint-Lac1

Within 48 h after transfection of pcDNA3.1^(−)^-Paint-Lac1, HeLa cells were washed with D-phosphate-buffered saline (D-PBS, Nacalai Tesque, #14249-95), collected by trypsinization, and pelleted by centrifugation. Cells were resuspended in HBSS supplemented with 10 mM HEPES, or in HBSS supplemented with 10 mM HEPES and 10 mM l-lactate. Samples were passed through a 35-μm cell strainer (Falcon, #352235) before running the samples. When required, whole-sample illumination with 405-nm light for 30 seconds was applied using the LED Flood Controller. Flow cytometry analysis was performed on an SH800 (Sony) with a 488-nm excitation laser and a 600/60-nm emission filter. Data were analyzed using FlowJo v10 software.

#### Imaging of Paint-Lac1-EGFP in HeLa cells

The pcDNA3.1^(−)^ plasmid encoding Paint-Lac1-EGFP was synthesized and used to transfect HeLa cells as described above for Paint-Lac1 imaging. Cells were pre-treated with iodoacetic acid, nigericin and rotenone and incubated in buffers containing l-lactate gradients. Whole-dish illumination with 405-nm light for 20 seconds (5 seconds on and 5 seconds off, 4 times) was applied using the LED Flood Controller. Fluorescence images were acquired before and after photoactivation on an IX83 wide-field microscope (Olympus) equipped with a pE-300 LED light source (CoolLED), a 20× objective, an ImagEM X2 EM-CCD camera (Hamamatsu), and cellSens software. Appropriate filter sets were used for the EGFP and Paint-Lac1 channels: the 510-nm channel was imaged using a 470/20 nm excitation filter, a 490 nm dichroic mirror, and a 518/45 nm emission filter; 560-nm channel was imaged using 438/24 nm excitation filter, a 458 nm dichroic mirror, and a 542/27 nm emission filter. Image acquisition was controlled using cellSens software, and all fluorescence data were analyzed in ImageJ software.

### QUANTIFICATION AND STATISTICAL ANALYSIS

Level of replication, statistical test, sample sizes, and error bars are stated in the figure legends.

The sensitivity of GR-iLACCO is reported as Δ*F*/*F*_min_ [= *F*^l-lactate(−)^ − *F*^l-lactate(+)^) / *F*^l-lactate(+)^], the sensitivity of Paint-Lac is reported as 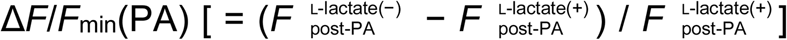, and the sensitivity of Paint-Lac1-EGFP is reported as Δ*R*′/*R*′_min_, where *R*′ = *R* – *R*_0_, *R* = *F*_560_ / *F*_510_ and *R*_0_ is the pre-activation ratio.

GraphPad Prism 10 was used for curve fittings. The sensitivity of the sensors as a function of l-lactate concentration was fitted to the Hill equation [*f*(*x*) = bottom + (top – bottom) / (1 + (EC_50_ / *x*)*^n^*)] to determine the EC_50_, where *n* is the Hill coefficient. For pH titration, pH buffer (30 mM MOPS, 30 mM trisodium citrate, 30 mM sodium borate, 100 mM KCl, and either no l-lactate or 10 mM l-lactate, with pH ranging from 5 to 10) was used to dilute protein solutions. Fluorescence intensities as a function of pH were then fitted to a sigmoidal function [*f*(pH) = bottom + (top – bottom) / (1 + 10^(p*K*_a_ − pH))] to determine the p*K*_a_.

## Notes

### Competing Interest Statement

The authors have declared no competing interest.

