## Supplementary information for Paint-Lac for "Paint-Lac: A photoactivatable integrator for l-lactate"

**A**

| Variant name | N-linker | C-linker | $\Delta F/F_{min}$ |
|---|---|---|---|
| GR-iLACCO0.1 | | KKTLKWMPSWGGSVPPVFS | 0.23 |
| GR-iLACCO0.1a | | KKTLKWMPSWGSVPPVFS | 0.17 |
| GR-iLACCO0.1b | SRQRMYRGGSEKIYVR | KKTLKWMPSWGVPPVFS | 0.23 |
| GR-iLACCO0.1c | | KKTLKWMPSWSVPPVFS | 0.13 |
| GR-iLACCO0.1d | | KKTLKWMPSWVPPVFS | **0.82** |

**B**

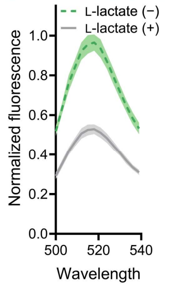


**C**

| Variant name | N-linker | C-linker | $\Delta F/F_{min}$ |
|---|---|---|---|
| GR-iLACCO0.1e | | KKTLKWMPSWGSPVFS | **1.25** |
| GR-iLACCO0.1f | SRQRMYRGGSEKIYVR | KKTLKWMPSWGPVFS | 0.57 |
| GR-iLACCO0.1g | | KKTLKWMPSWSPVFS | 0.49 |
| GR-iLACCO0.1h | | KKTLKWMPSWPVFS | 0.20 |

**D**

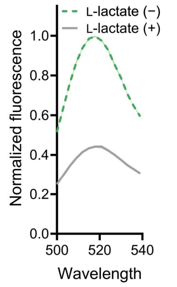


**E**

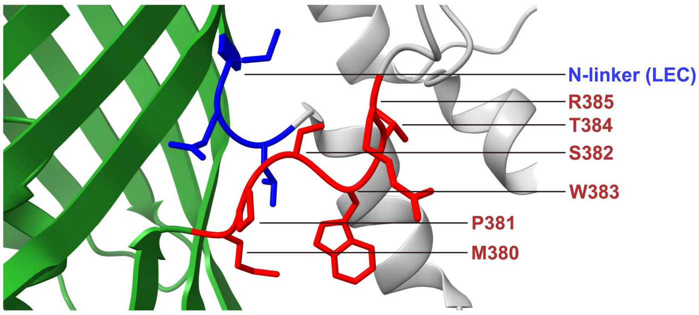


**F**

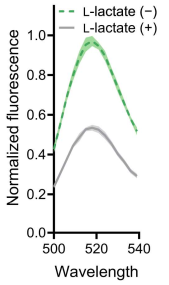


**Figure S1**. **Linker optimization from GR-iLACCO0.1 to GR-iLACCO0.3.** (**A**, **C**) GR-iLACCO0.1 variants showing their corresponding C-linker sequences and $\Delta F/F_{min}$ values in the green state measured using crude protein extract by B-PER. (**B**, **D**) Emission spectra (excitation at 500 nm) of GR-iLACCO0.1d and GR-iLACCO0.1e in the presence or absence of L-lactate, measured in crude protein extract by B-PER. These variants were selected as templates for the next optimization step. (**E**) Linker amino-acid residues illustrated on the crystal structure of calcium-free CaMPARI v0.2[1] (PDB ID: 4oy4), with N-linker highlighted in blue and C-linker-related residues in red. The cpmEos2 domain is shown in green, and the CaM and M13 domains are shown in gray. (**F**) Emission spectra (excitation at 500 nm) of GR-iLACCO0.3 in the presence or absence of L-lactate, measured using Ni-NTA-purified protein.

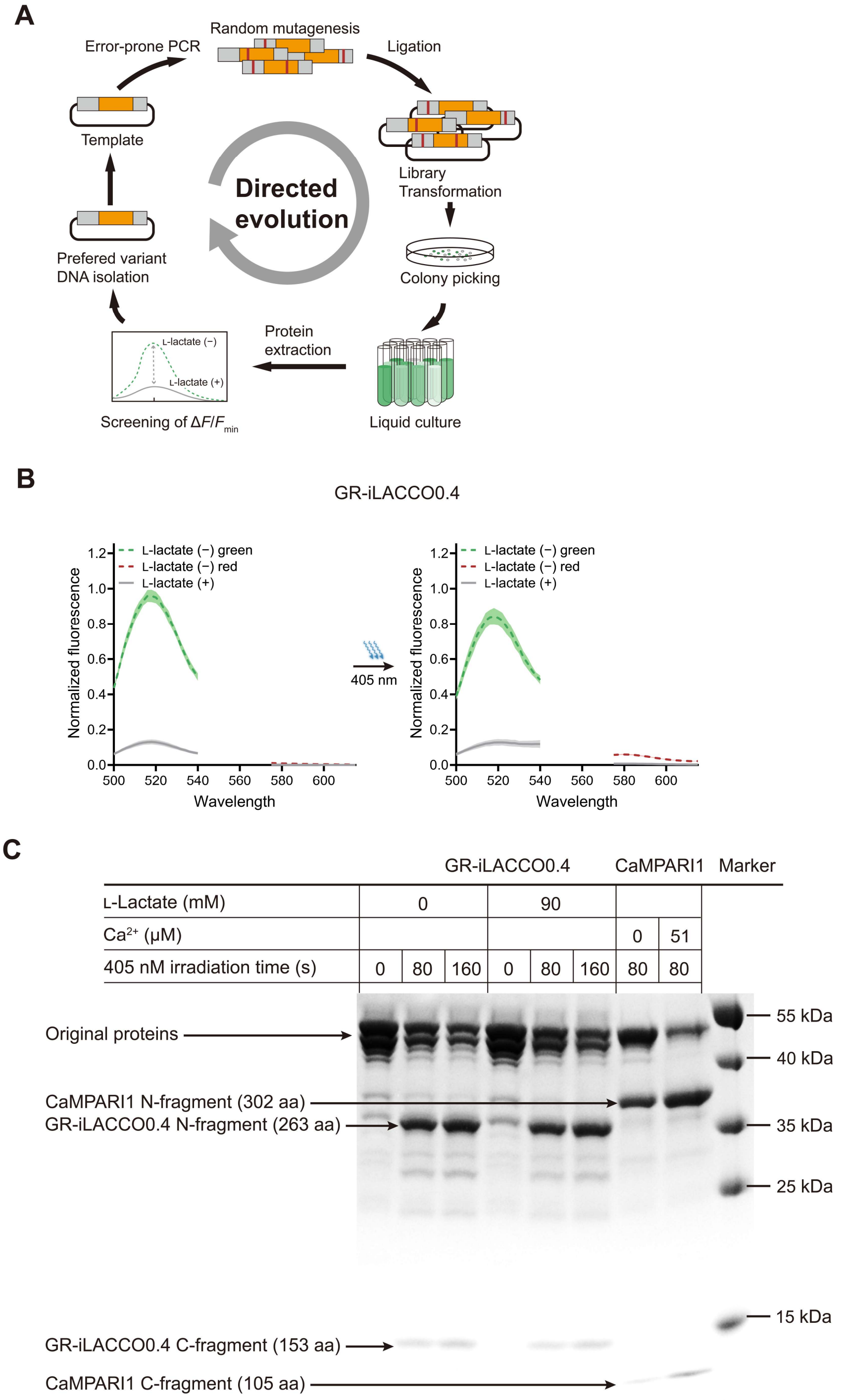


**Figure S2**. **Development and characterization of GR-iLACCO0.4.** (**A**) Directed evolution protocol for the development of GR-iLACCO. After error-prone PCR and construction of the plasmid library, bright colonies on agar plate were picked for the following culture. The $\Delta F/F_{min}$ of the green state was determined using crude protein extract by B-PER. Gene encoding the variant with the highest $\Delta F/F_{min}$ was used as template for the next round. (**B**) Emission spectra (excitation at 500 nm for green channel and at 560 nm for red channel) of GR-iLACCO0.4 before and after irradiation in the presence or absence of L-lactate, measured using Ni-NTA-purified protein. (**C**) SDS-PAGE comparing GR-iLACCO0.4 and CaMPARI1 in terms of their light-induced cleavage.

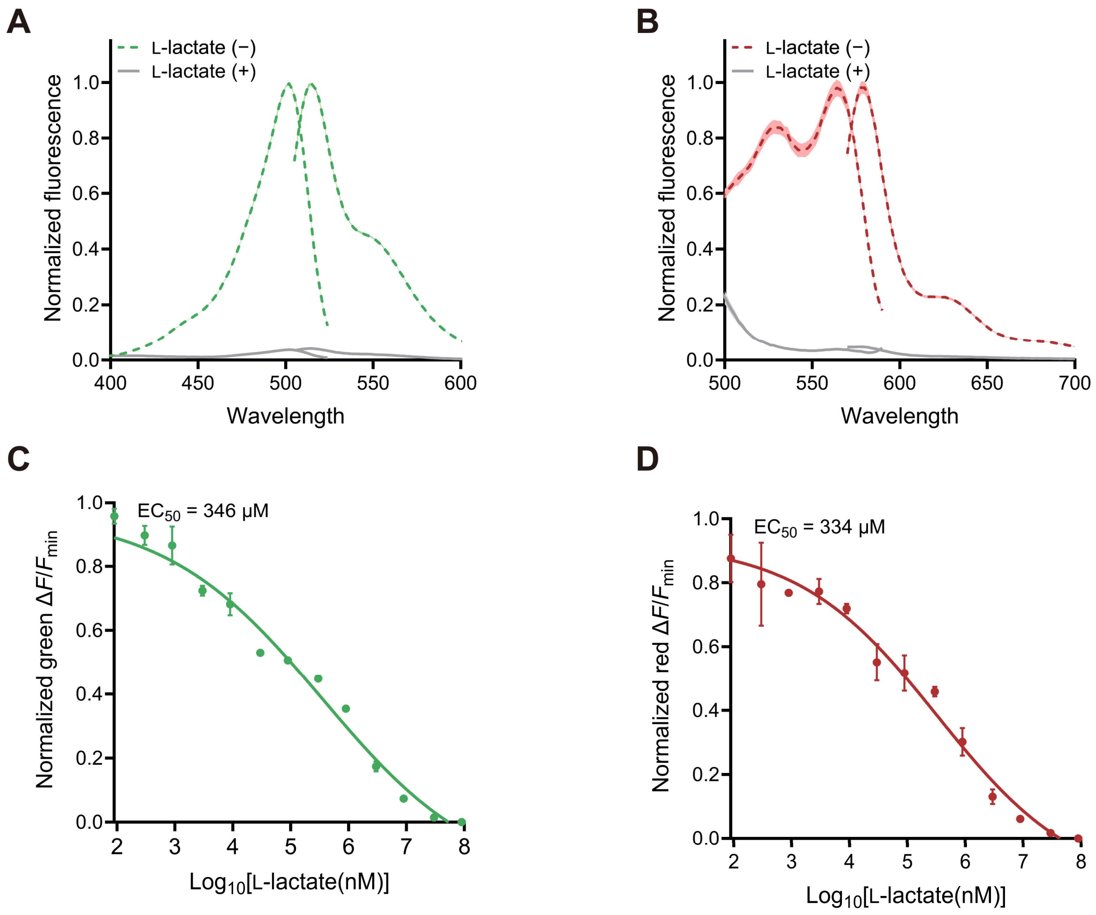


**Figure S3**. **Characterization of GR-iLACCO latest variant, GR-iLACCO1.** (**A, B**) Excitation (emission at 520 nm for green state and at 580 nm for red state) and emission (excitation at 500 nm for green state and at 560 nm for red state) spectra of green-state and red-state GR-iLACCO1, measured using Ni-NTA-purified protein. (**C, D**) Dose–response curves of GR-iLACCO1 green and red states for L-lactate, measured using Ni-NTA-purified protein. ($n$ = 3 technical replicates, mean ± s.d.).

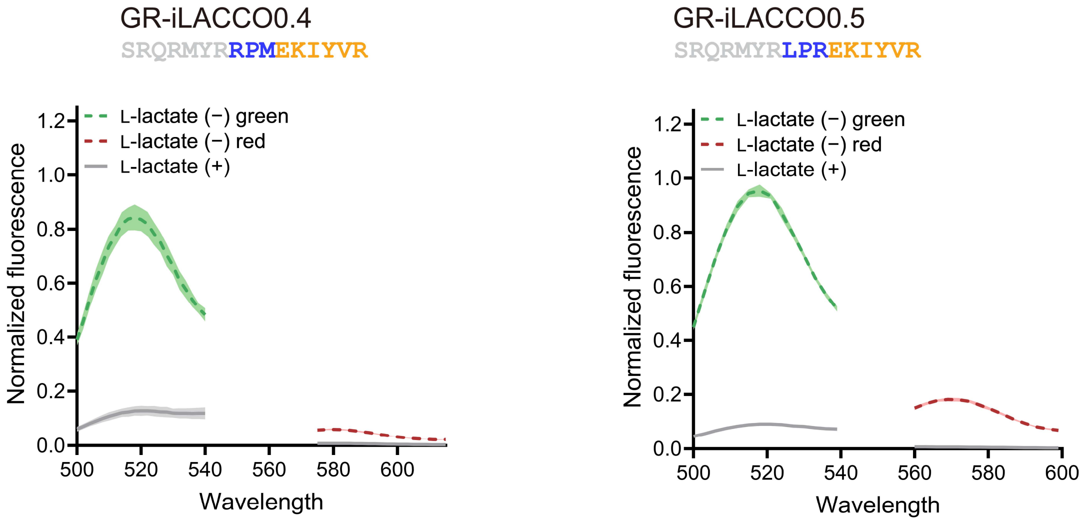


31 **Figure S4**. **Second-round N-linker optimization of GR-iLACCO.** Emission spectra (excitation at 500 nm
32 for green channel and at 560 nm for red channel) of GR-iLACCO0.4 and GR-iLACCO0.5 upon irradiation
33 in the presence or absence of L-lactate, measured in crude protein extract by B-PER.

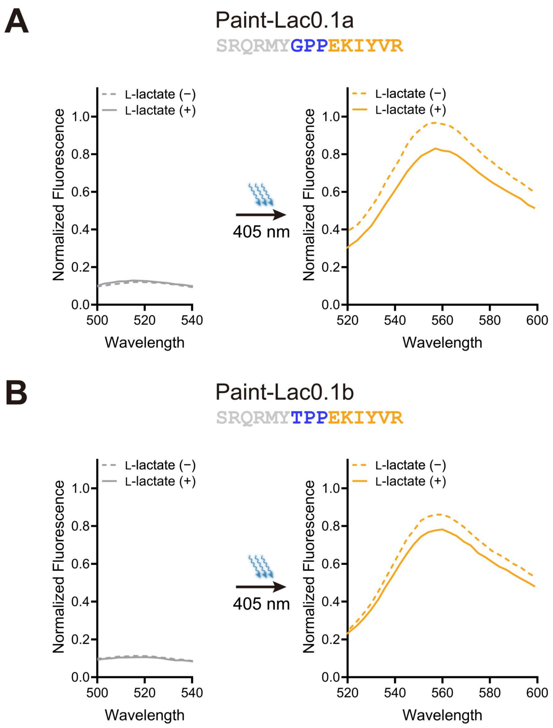


**Figure S5**. **Photoactivation of Paint-Lac prototypes.** (**A**, **B**) Emission spectra (excitation at 450 nm both the dim and bright orange states) of Paint-Lac1a/b before and after irradiation in the presence (90 mM) and absence of L-lactate, measured using crude protein extracts. The corresponding N-linker sequences for each prototype are shown.

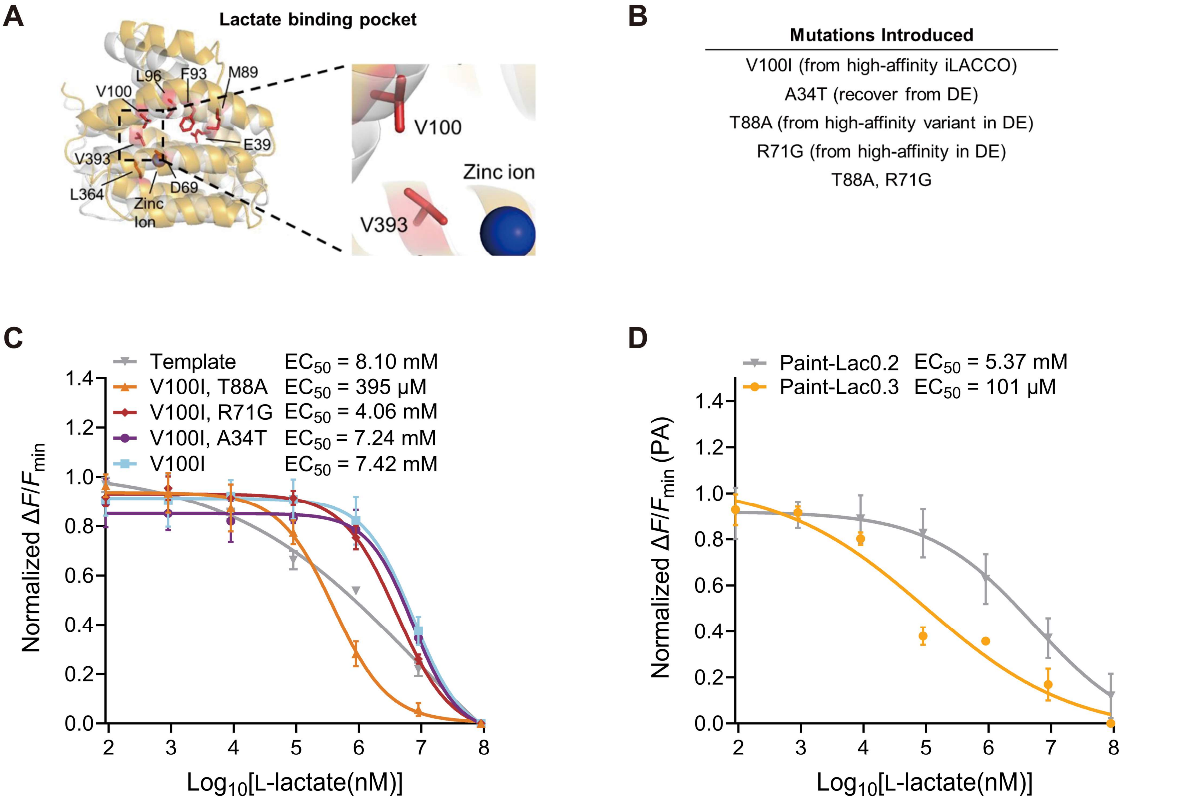


**Figure S6**. **Tuning of GR-iLACCO affinity through structure-guided mutagenesis.** (**A**) Structural model of the L-lactate-binding pocket showing residues selected for affinity tuning of iLACCO. Key residues are highlighted in red, and a zinc ion is shown in the binding site. Adapted from ref. 15 and used with permission. (**B**) Mutations introduced include V100I (from iLACCO1), A34T (reversion of an evolution-derived substitution), and T88A and R71G (from a high-affinity variant). (**C**) Dose–response curves for GR-iLACCO variants containing combinations of affinity-enhancing mutations, measured using Ni-NTA-purified protein. “Template” refers to GR-iLACCO0.5 (*n* = 3 technical replicates, mean ± s.d.). (**D**) Dose–response curves for Paint-Lac0.2 and Paint-Lac0.3, measured using Ni-NTA-purified protein (*n* = 3 technical replicates, mean ± s.d.).

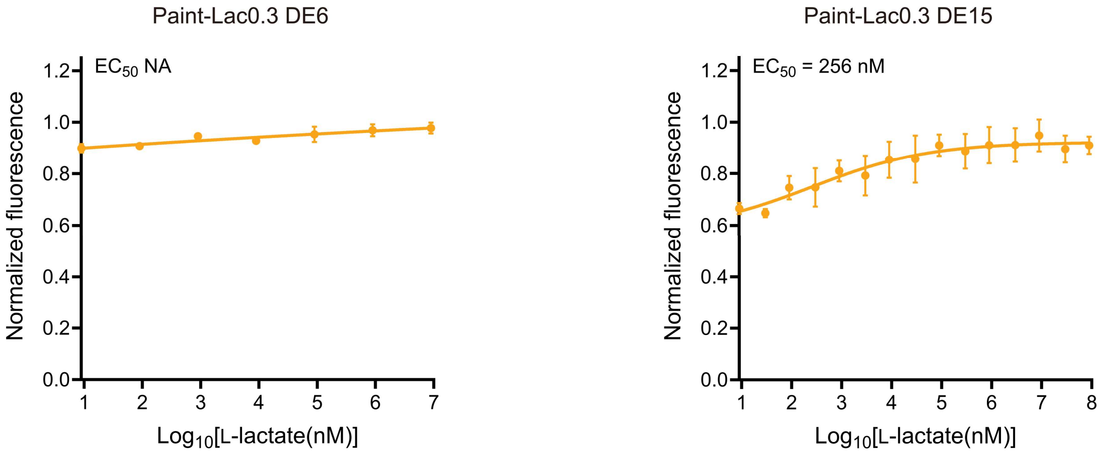


**Figure S7**. **Signal stability of the bright orange state during directed evolution.** Dose–response curves of Paint-Lac0.3-DE6 and -DE15 variant (after 6 and 15 rounds of directed evolution, respectively) bright orange states for L-lactate, measured using Ni-NTA-purified protein (*n* = 3 technical replicates, mean ± s.d.).

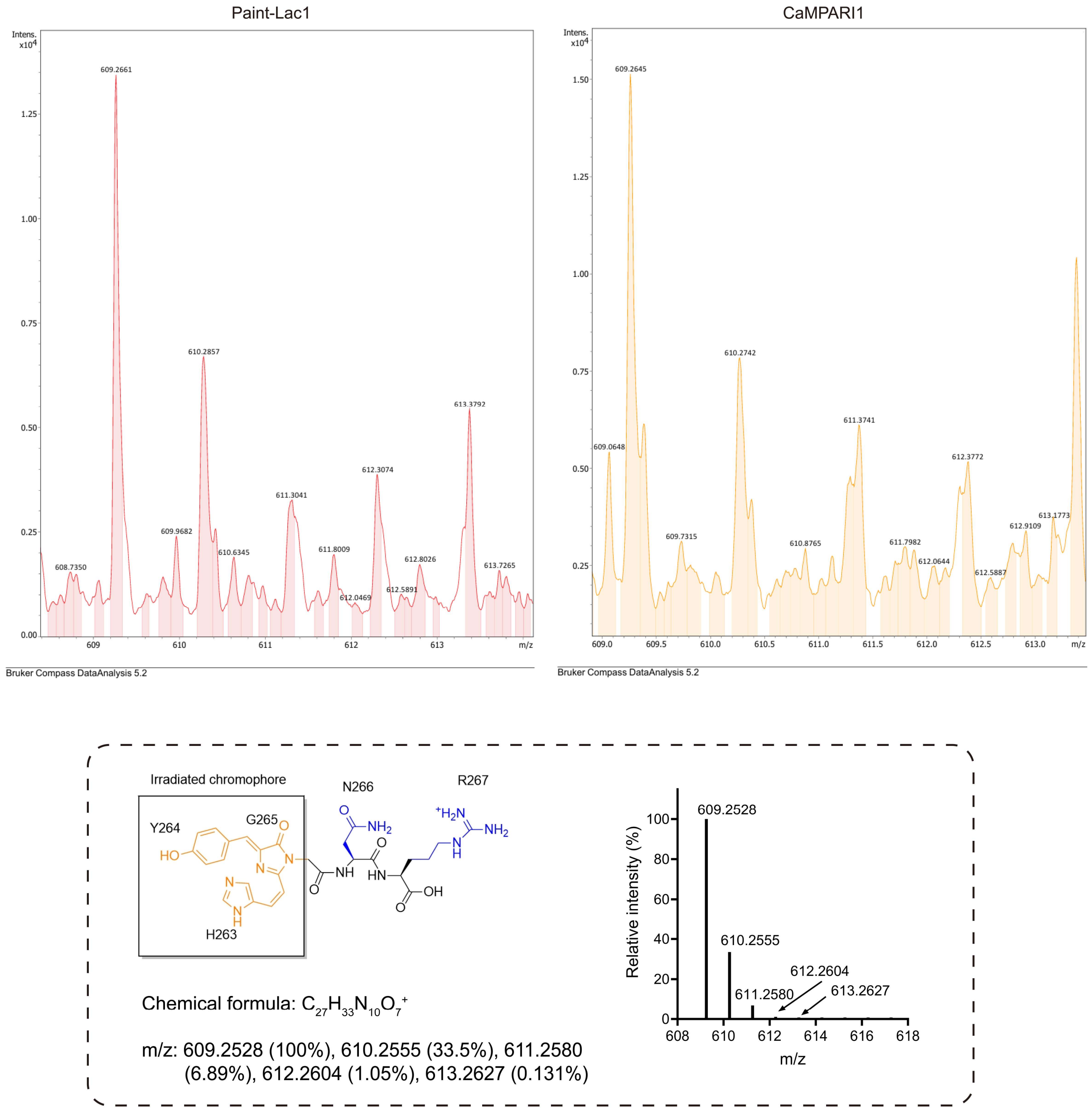


**Figure S8. Mass spectra comparing the chromophore-containing peptides generated from bright orange Paint-Lac1 and red CaMPARI1.** The expected fragment was shown at the bottom. Amino residues are numbered as in Paint-Lac1. The bright orange state of HYG-form chromophore is shown in orange; downstream residues are shown in blue; and the peptide backbone is shown in black. The monoisotopic mass and isotopic pattern were generated using ChemCalc, an online isotopic distribution calculator[2].

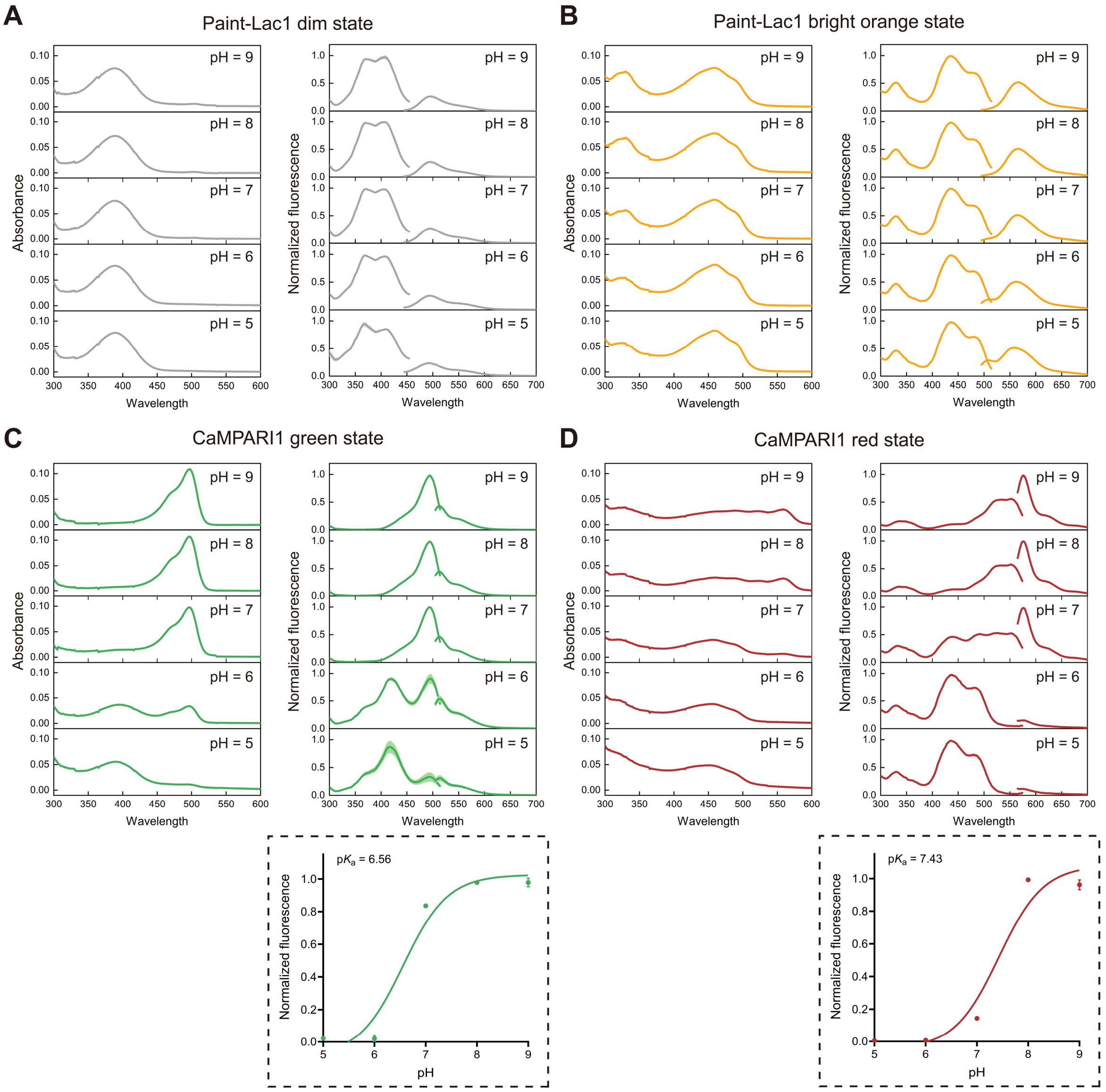


**Figure S9**. **Photophysical properties of Paint-Lac1 and CaMPARI1 across pH 5–9.** (**A**, **B**) Absorbance, excitation (emission at 500 nm for the dim state and 560 for the bright orange state), and emission (excitation at 400 for the dim state and 450 for the bright orange state) spectra of Paint-Lac1 dim and bright orange states measured across pH 5–9. (**C**, **D**) Absorbance, excitation (emission at 520 nm for the green state and 580 for the red state), and emission (excitation at 500 nm for green state and 560 for red state) spectra of CaMPARI1 green and red states measured across pH 5–9. Corresponding pH titration curves for the green and red states are shown ($n$ = 3 technical replicates, mean ± s.d.).

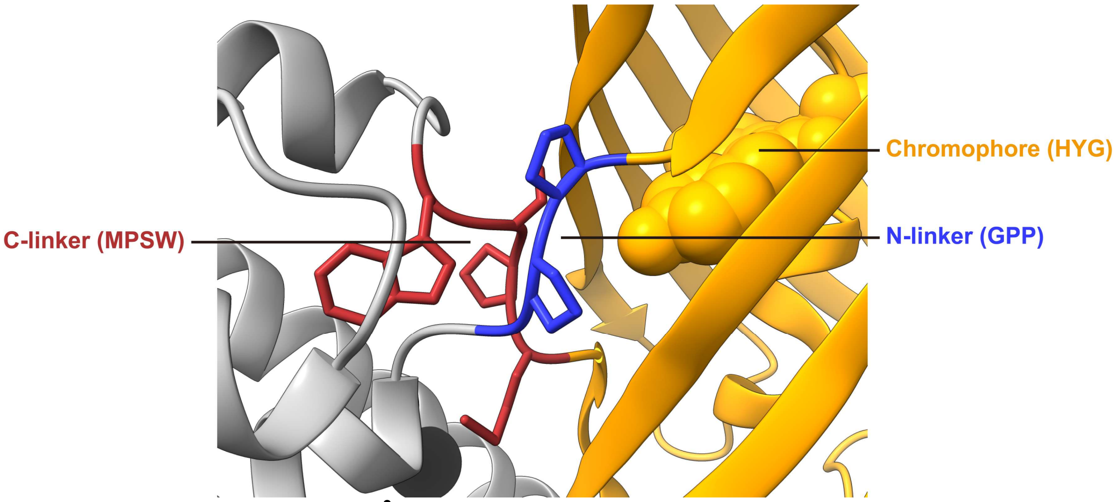


**Figure S10. AlphaFold3 model[3] highlights the N- and C-linker regions and chromophore environment in Paint-Lac1.** The HYG-formed chromophore, shown in orange, is from the CaMPARI v0.2 crystal structure[1] (PDB ID: 4oy4) that is superimposed with Paint-Lac1 model.

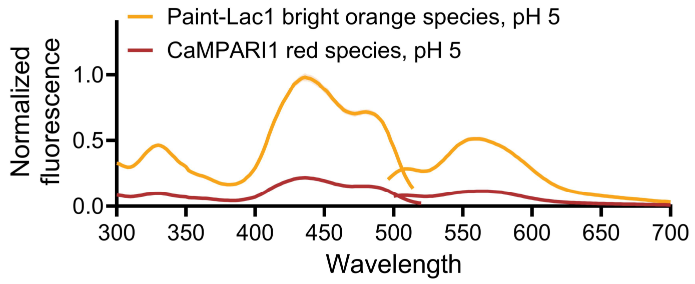


**Figure S11. Excitation (emission at 560 nm) and emission (excitation at 450 nm) spectra of bright orange Paint-Lac1 and red CaMPARI1 measured at pH 5.**

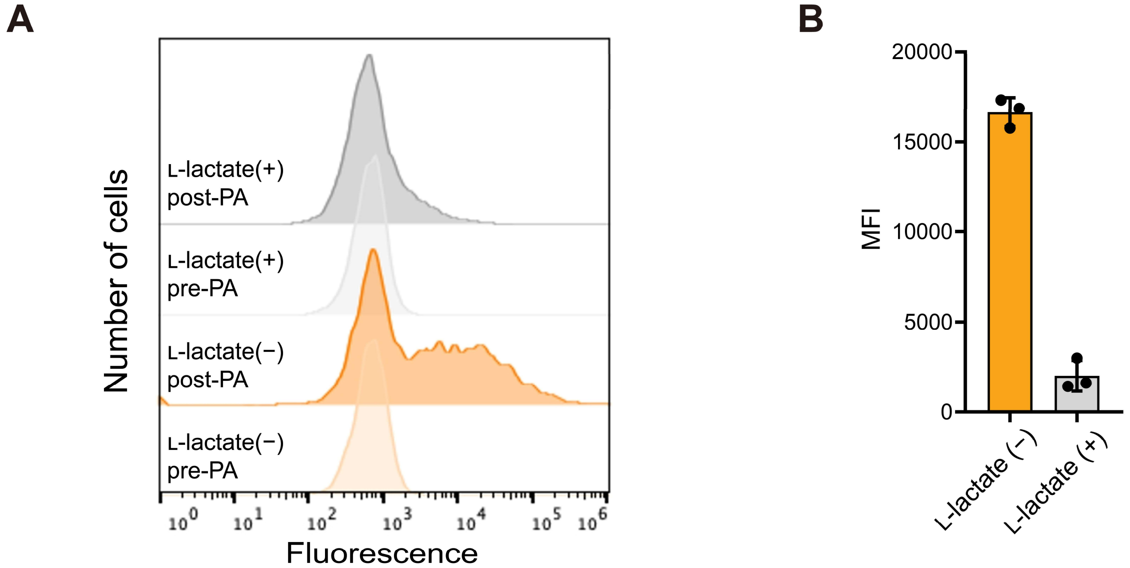


**Figure S12. Flow cytometry analysis of HeLa cells expressing Paint-Lac1.** (**A**) Representative flow cytometry histograms. (**B**) Mean fluorescent intensity (MFI) values of cells irradiated in the presence or absence of L-lactate ($n$ = 3 technical replicates, mean ± s.d., and approximately 1.0×$10^5$ cells were analyzed for each independent experiment.).

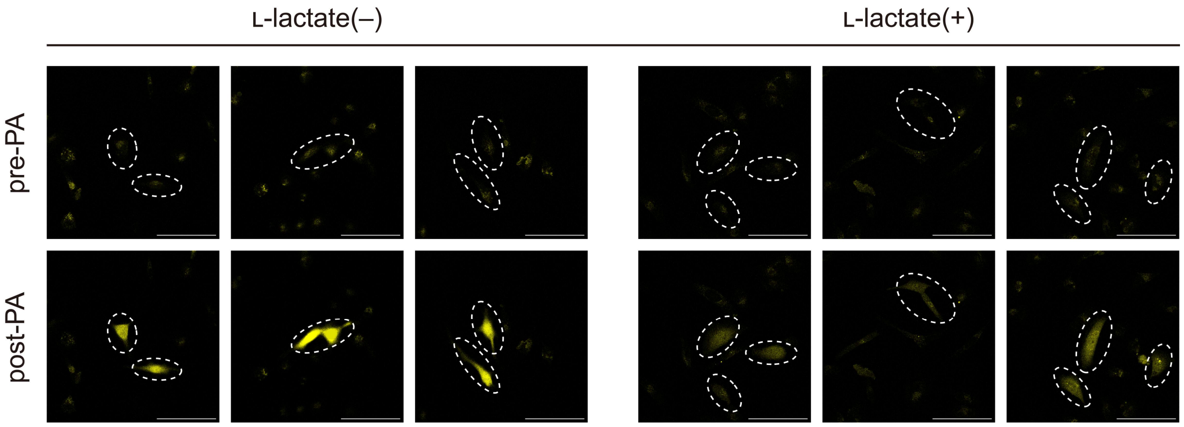


**Figure S13**. **Confocal microscopy of Paint-Lac1 in cells.** Images of HeLa cells expressing Paint-Lac1. Cell-selected irradiation was applied with 405-nm light in the absence or presence of 5 mM L-lactate. White dash circles represent the areas of irradiation. Scale bars, 100 μm.

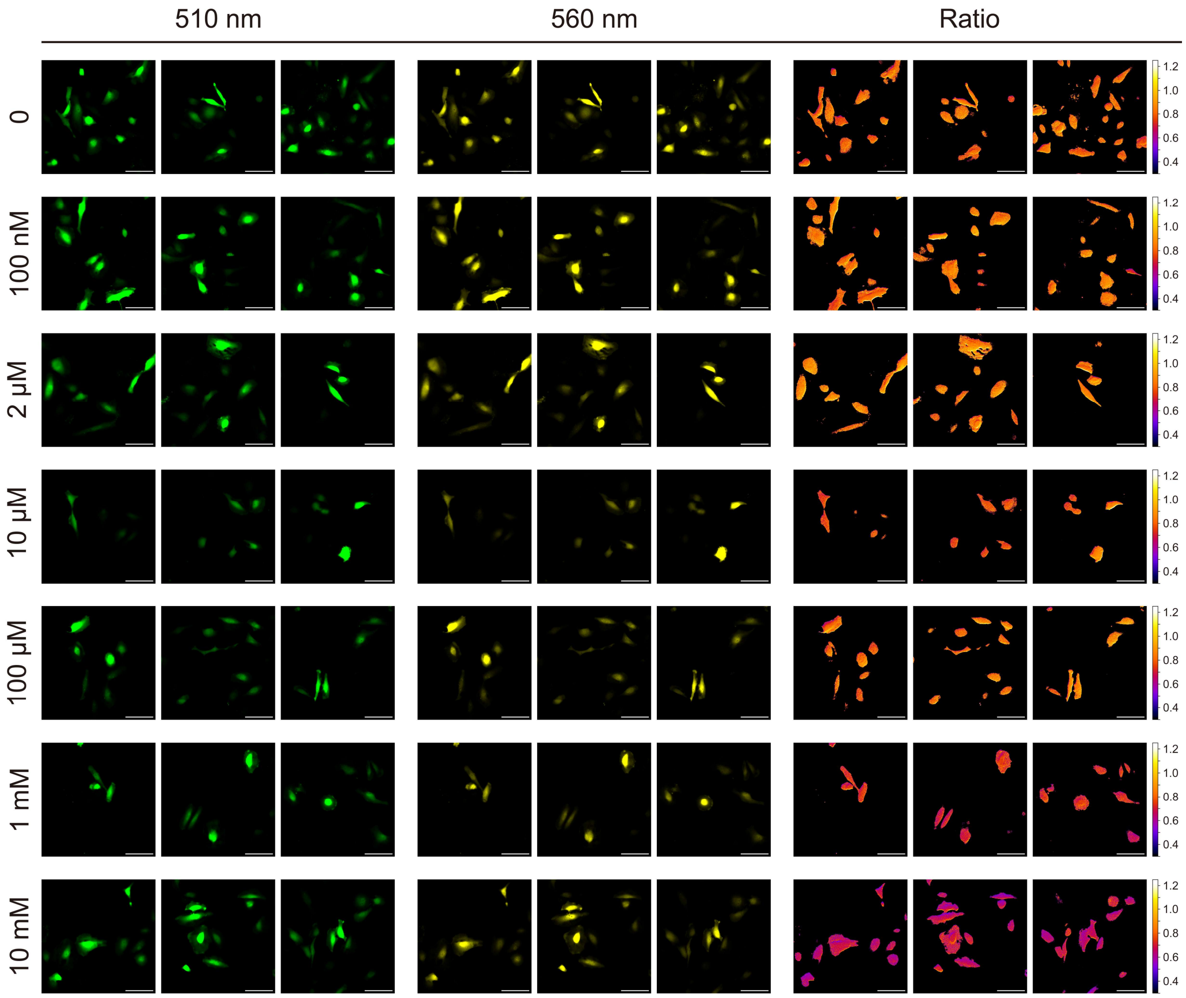


**Figure S14**. **Fluorescence microscopy of Paint-Lac1-EGFP in cells.** Images of HeLa cells expressing Paint-Lac1-EGFP and irradiated with 405-nm light in L-lactate gradient. Scale bars, 50 μm.

| Paint-Lac1 #s | 1 | 2 | 3 | 4 | 5 | 6 | 7 | 8 | 9 | 10 | 11 | 12 | 13 | 14 | 15 | 16 | 17 | 18 | 19 | 20 | 21 | 22 | 23 | 24 | 25 | 26 | 27 | 28 | 29 | 30 | 31 | 32 | 33 | 34 | 35 | 36 | 37 | 38 | 39 | 40 |
|---|---|---|---|---|---|---|---|---|---|---|---|---|---|---|---|---|---|---|---|---|---|---|---|---|---|---|---|---|---|---|---|---|---|---|---|---|---|---|---|---|
| Paint-Lac1 | M | E | Q | N | I | A | Q | P | L | K | T | H | M | A | D | D | P | N | Y | A | Y | D | I | L | E | A | R | Y | A | I | E | A | S | A | A | W | H | A | A | M |
| GR-iLACCO0.4 | M | E | Q | N | I | V | Q | P | L | K | T | H | M | A | D | D | P | D | Y | T | F | D | I | L | E | A | R | Y | A | I | E | A | S | A | A | W | H | A | A | M |
| LldR | M | E | Q | N | I | V | Q | P | L | K | T | H | M | A | D | D | P | D | Y | T | F | D | I | L | E | A | R | Y | A | I | E | A | S | T | A | W | H | A | A | M |
| cpmEos2 | - | - | - | - | - | - | - | - | - | - | - | - | - | - | - | - | - | - | - | - | - | - | - | - | - | - | - | - | - | - | - | - | - | - | - | - | - | - | - | - |

| | 41 | 42 | 43 | 44 | 45 | 46 | 47 | 48 | 49 | 50 | 51 | 52 | 53 | 54 | 55 | 56 | 57 | 58 | 59 | 60 | 61 | 62 | 63 | 64 | 65 | 66 | 67 | 68 | 69 | 70 | 71 | 72 | 73 | 74 | 75 | 76 | 77 | 78 | 79 | 80 |
|---|---|---|---|---|---|---|---|---|---|---|---|---|---|---|---|---|---|---|---|---|---|---|---|---|---|---|---|---|---|---|---|---|---|---|---|---|---|---|---|---|
| Paint-Lac1 | R | A | T | P | G | D | K | E | K | I | R | L | C | F | E | A | T | L | S | E | D | P | D | I | A | S | Q | A | D | V | R | F | H | L | A | I | A | E | A | S |
| GR-iLACCO0.4 | R | A | T | P | G | D | K | E | K | I | R | L | C | F | E | A | T | L | S | E | D | P | D | I | A | S | Q | A | D | V | R | F | H | L | A | I | A | E | A | S |
| LldR | R | A | T | P | G | D | K | E | K | I | Q | L | C | F | E | A | T | L | S | E | D | P | D | I | A | S | Q | A | D | V | R | F | H | L | A | I | A | E | A | S |
| cpmEos2 | - | - | - | - | - | - | - | - | - | - | - | - | - | - | - | - | - | - | - | - | - | - | - | - | - | - | - | - | - | - | - | - | - | - | - | - | - | - | - | - |

| | 81 | 82 | 83 | 84 | 85 | 86 | 87 | 88 | 89 | 90 | 91 | 92 | 93 | 94 | 95 | 96 | 97 | 98 | 99 | 100 | 101 | 102 | 103 | 104 | 105 | 106 | 107 | 108 | - | 109 | 110 | 111 | 112 | 113 | 114 | 115 | 116 | 117 | 118 | 119 |
|---|---|---|---|---|---|---|---|---|---|---|---|---|---|---|---|---|---|---|---|---|---|---|---|---|---|---|---|---|---|---|---|---|---|---|---|---|---|---|---|---|
| Paint-Lac1 | H | N | I | A | L | L | Q | A | M | R | G | Y | F | D | A | L | Q | S | S | I | R | H | S | R | Q | R | M | Y | - | G | P | P | E | R | I | Y | V | R | D | G |
| GR-iLACCO0.4 | H | N | I | A | L | L | Q | T | M | R | G | Y | F | D | V | L | Q | S | S | V | R | H | S | R | Q | R | M | Y | R | L | P | R | E | R | I | Y | V | R | D | G |
| LldR | H | N | I | V | L | L | Q | T | M | R | G | Y | F | D | V | L | Q | S | S | V | R | H | S | R | Q | R | M | Y | R | - | - | - | - | - | - | - | - | - | - | - |
| cpmEos2 | - | - | - | - | - | - | - | - | - | - | - | - | - | - | - | - | - | - | - | - | - | - | - | - | - | - | - | - | - | - | - | - | E | K | I | Y | V | R | D | G |

| | 120 | 121 | 122 | 123 | 124 | 125 | 126 | 127 | 128 | 129 | 130 | 131 | 132 | 133 | 134 | 135 | 136 | 137 | 138 | 139 | 140 | 141 | 142 | 143 | 144 | 145 | 146 | 147 | 148 | 149 | 150 | 151 | 152 | 153 | 154 | 155 | 156 | 157 | 158 | 159 |
|---|---|---|---|---|---|---|---|---|---|---|---|---|---|---|---|---|---|---|---|---|---|---|---|---|---|---|---|---|---|---|---|---|---|---|---|---|---|---|---|---|
| Paint-Lac1 | V | L | T | G | D | I | H | M | I | L | L | L | E | G | N | T | L | Y | R | C | D | F | R | T | A | Y | K | A | K | E | K | G | V | K | L | P | D | E | H | F |
| GR-iLACCO0.4 | V | L | T | G | D | I | H | M | I | L | L | L | E | G | N | T | H | Y | R | C | D | F | R | T | A | Y | K | A | K | E | K | G | V | K | L | P | G | V | H | F |
| LldR | - | - | - | - | - | - | - | - | - | - | - | - | - | - | - | - | - | - | - | - | - | - | - | - | - | - | - | - | - | - | - | - | - | - | - | - | - | - | - | - |
| cpmEos2 | V | L | T | G | D | I | H | M | I | L | L | L | E | G | N | A | H | Y | R | C | D | F | R | T | T | Y | K | A | K | E | K | G | V | K | L | P | G | V | H | F |

| | 160 | 161 | 162 | 163 | 164 | 165 | 166 | 167 | 168 | 169 | 170 | 171 | 172 | 173 | 174 | 175 | 176 | 177 | 178 | 179 | 180 | 181 | 182 | 183 | 184 | 185 | 186 | 187 | 188 | 189 | 190 | 191 | 192 | 193 | 194 | 195 | 196 | 197 | 198 | 199 |
|---|---|---|---|---|---|---|---|---|---|---|---|---|---|---|---|---|---|---|---|---|---|---|---|---|---|---|---|---|---|---|---|---|---|---|---|---|---|---|---|---|
| Paint-Lac1 | V | D | H | C | I | E | I | L | S | H | D | E | D | Y | N | K | V | K | L | Y | E | Y | A | V | A | H | P | G | L | P | D | N | A | G | R | G | G | T | G | G |
| GR-iLACCO0.4 | V | D | H | C | I | E | I | L | S | H | D | K | D | Y | N | K | V | K | L | Y | E | Y | A | V | A | H | P | G | L | P | D | N | A | G | R | G | G | T | G | G |
| LldR | - | - | - | - | - | - | - | - | - | - | - | - | - | - | - | - | - | - | - | - | - | - | - | - | - | - | - | - | - | - | - | - | - | - | - | - | - | - | - | - |
| cpmEos2 | V | D | H | C | I | E | I | L | S | H | D | K | D | Y | N | K | V | K | L | Y | E | Y | A | V | A | H | S | G | L | P | D | N | A | R | R | G | G | T | G | G |

| | 200 | 201 | 202 | 203 | 204 | 205 | 206 | 207 | 208 | 209 | 210 | 211 | 212 | 213 | 214 | 215 | 216 | 217 | 218 | 219 | 220 | 221 | 222 | 223 | 224 | 225 | 226 | 227 | 228 | 229 | 230 | 231 | 232 | 233 | 234 | 235 | 236 | 237 | 238 | 239 |
|---|---|---|---|---|---|---|---|---|---|---|---|---|---|---|---|---|---|---|---|---|---|---|---|---|---|---|---|---|---|---|---|---|---|---|---|---|---|---|---|---|
| Paint-Lac1 | S | M | V | S | A | I | E | P | D | M | K | N | K | L | R | M | E | G | N | V | N | G | H | H | F | V | I | D | G | V | G | T | G | K | P | Y | E | G | S | Q |
| GR-iLACCO0.4 | S | M | V | S | A | I | K | P | D | M | K | I | K | L | R | M | E | G | N | V | N | G | H | H | F | V | I | D | G | D | G | T | G | K | P | Y | E | G | N | Q |
| LldR | - | - | - | - | - | - | - | - | - | - | - | - | - | - | - | - | - | - | - | - | - | - | - | - | - | - | - | - | - | - | - | - | - | - | - | - | - | - | - | - |
| cpmEos2 | S | M | V | S | A | I | K | P | D | M | K | I | K | L | R | M | E | G | N | V | N | G | H | H | F | V | I | D | G | D | G | T | G | K | P | Y | E | G | K | Q |

| | 240 | 241 | 242 | 243 | 244 | 245 | 246 | 247 | 248 | 249 | 250 | 251 | 252 | 253 | 254 | 255 | 256 | 257 | 258 | 259 | 260 | 261 | 262 | 263 | 264 | 265 | 266 | 267 | 268 | 269 | 270 | 271 | 272 | 273 | 274 | 275 | 276 | 277 | 278 | 279 |
|---|---|---|---|---|---|---|---|---|---|---|---|---|---|---|---|---|---|---|---|---|---|---|---|---|---|---|---|---|---|---|---|---|---|---|---|---|---|---|---|---|
| Paint-Lac1 | T | M | D | L | E | V | K | E | G | G | P | L | P | F | A | F | D | I | L | T | T | A | F | H | Y | G | N | R | V | F | V | K | Y | P | D | N | I | Q | D | Y |
| GR-iLACCO0.4 | T | M | D | L | E | V | K | E | G | G | P | L | P | F | A | F | D | I | L | T | T | A | F | H | Y | G | N | R | V | F | V | K | Y | P | D | N | I | Q | D | Y |
| LldR | - | - | - | - | - | - | - | - | - | - | - | - | - | - | - | - | - | - | - | - | - | - | - | - | - | - | - | - | - | - | - | - | - | - | - | - | - | - | - | - |
| cpmEos2 | T | M | D | L | E | V | K | E | G | G | P | L | P | F | A | F | D | I | L | T | T | A | F | H | Y | G | N | R | V | F | V | K | Y | P | D | N | I | Q | D | Y |

| | 280 | 281 | 282 | 283 | 284 | 285 | 286 | 287 | 288 | 289 | 290 | 291 | 292 | 293 | 294 | 295 | 296 | 297 | 298 | 299 | 300 | 301 | 302 | 303 | 304 | 305 | 306 | 307 | 308 | 309 | 310 | 311 | 312 | 313 | 314 | 315 | 316 | 317 | 318 | 319 |
|---|---|---|---|---|---|---|---|---|---|---|---|---|---|---|---|---|---|---|---|---|---|---|---|---|---|---|---|---|---|---|---|---|---|---|---|---|---|---|---|---|
| Paint-Lac1 | F | K | Q | S | F | P | K | G | Y | S | W | E | R | S | M | T | F | E | D | G | G | I | C | H | A | R | N | D | I | T | M | E | G | D | T | F | Y | N | K | V |
| GR-iLACCO0.4 | F | K | Q | S | F | P | K | G | Y | S | W | E | R | S | M | T | F | E | D | G | G | I | C | H | A | R | N | D | I | T | M | E | G | D | T | F | Y | N | K | V |
| LldR | - | - | - | - | - | - | - | - | - | - | - | - | - | - | - | - | - | - | - | - | - | - | - | - | - | - | - | - | - | - | - | - | - | - | - | - | - | - | - | - |
| cpmEos2 | F | K | Q | S | F | P | K | G | Y | S | W | E | R | S | M | T | F | E | D | G | G | I | C | Y | A | R | N | D | I | T | M | E | G | D | T | F | Y | N | K | V |

| | 320 | 321 | 322 | 323 | 324 | 325 | 326 | 327 | 328 | 329 | 330 | 331 | 332 | 333 | 334 | 335 | 336 | 337 | 338 | 339 | 340 | 341 | 342 | 343 | 344 | 345 | 346 | 347 | 348 | 349 | 350 | 351 | 352 | 353 | 354 | 355 | 356 | 357 | 358 | 359 |
|---|---|---|---|---|---|---|---|---|---|---|---|---|---|---|---|---|---|---|---|---|---|---|---|---|---|---|---|---|---|---|---|---|---|---|---|---|---|---|---|---|
| Paint-Lac1 | R | F | Y | G | T | N | F | P | A | N | G | P | V | M | Q | K | K | T | L | K | W | M | P | S | W | G | S | P | V | F | S | Q | L | T | E | Q | H | Q | A | V |
| GR-iLACCO0.4 | R | F | Y | G | T | N | F | P | A | N | G | P | V | M | Q | K | K | T | L | K | W | M | P | S | W | G | S | P | V | F | S | Q | L | T | E | Q | H | Q | A | V |
| LldR | - | - | - | - | - | - | - | - | - | - | - | - | - | - | - | - | - | - | - | - | - | - | - | - | - | V | P | P | V | F | S | Q | L | T | E | Q | H | Q | A | V |
| cpmEos2 | R | F | Y | G | T | N | F | P | A | N | G | P | V | M | Q | K | K | T | L | K | W | M | P | S | W | - | - | - | - | - | - | - | - | - | - | - | - | - | - | - |

| | 360 | 361 | 362 | 363 | 364 | 365 | 366 | 367 | 368 | 369 | 370 | 371 | 372 | 373 | 374 | 375 | 376 | 377 | 378 | 379 | 380 | 381 | 382 | 383 | 384 | 385 | 386 | 387 | 388 | 389 | 390 | 391 | 392 | 393 | 394 | 395 | 396 | 397 | 398 | 399 |
|---|---|---|---|---|---|---|---|---|---|---|---|---|---|---|---|---|---|---|---|---|---|---|---|---|---|---|---|---|---|---|---|---|---|---|---|---|---|---|---|---|
| Paint-Lac1 | M | D | A | I | F | A | G | D | A | D | G | A | R | K | A | M | L | A | H | L | S | F | V | H | T | T | M | K | L | F | D | D | D | Q | A | R | H | A | R | I |
| GR-iLACCO0.4 | I | D | A | I | F | A | G | D | A | D | G | A | R | K | A | M | L | A | H | L | S | F | V | H | T | T | M | K | R | F | D | D | D | Q | A | R | H | A | R | I |
| LldR | I | D | A | I | F | A | G | D | A | D | G | A | R | K | A | M | L | A | H | L | S | F | V | H | T | T | M | K | R | F | D | D | D | H | A | R | H | A | R | I |
| cpmEos2 | - | - | - | - | - | - | - | - | - | - | - | - | - | - | - | - | - | - | - | - | - | - | - | - | - | - | - | - | - | - | - | - | - | - | - | - | - | - | - | - |

| | 400 | 401 | 402 | 403 | 404 | 405 | 406 | 407 | 408 | 409 | 410 | 411 | 412 | 413 | 414 | 415 | - |
|---|---|---|---|---|---|---|---|---|---|---|---|---|---|---|---|---|---|
| Paint-Lac1 | T | R | L | P | G | E | H | N | E | H | S | R | E | K | N | A | * |
| GR-iLACCO0.4 | T | R | L | P | G | E | H | N | E | H | S | R | E | K | N | A | * |
| LldR | T | R | L | P | G | E | H | N | E | H | S | R | E | K | N | A | * |
| cpmEos2 | - | - | - | - | - | - | - | - | - | - | - | - | - | - | - | - | - |

**Figure S15. Protein sequence alignment.** The LldR-LBD domain is indicated by a gray background and the cpmEos2 domain is indicated by an orange background. Mutated residues are shown by red background, mutations kept from GR-iLACCO0.4 are shown in black, affinity-tuning mutations are shown by magenta background, N- and C-linker are indicated by a blue background and a dark red background, respectively, corresponding to the colors in **Figure 2**. Chromophore residues are enclosed within a black line.

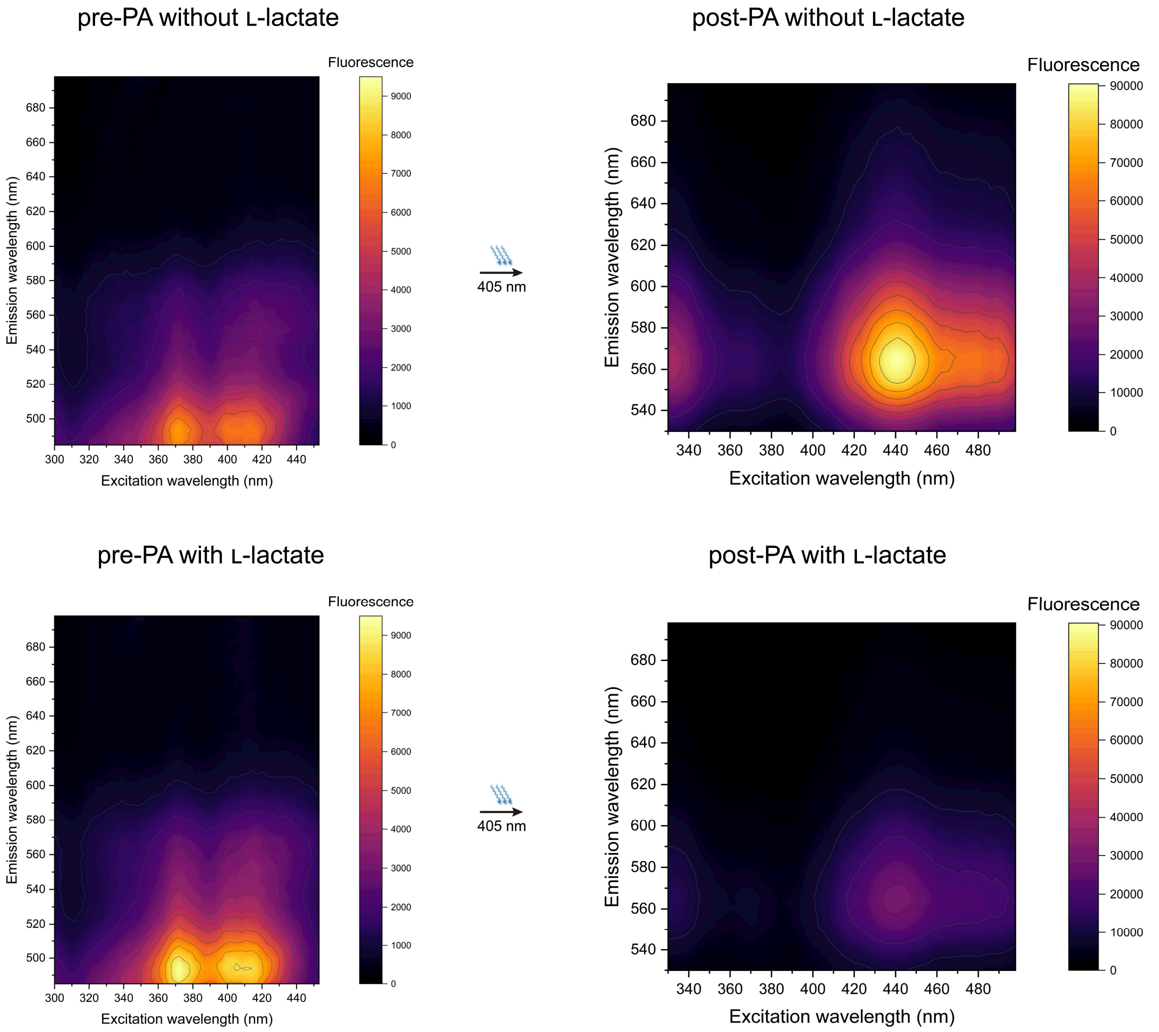


**Figure S16. Three-dimensional fluorescence spectra of Paint-Lac1**

### TABLE S1

| | Paint-Lac1 | | | | |
|---|---|---|---|---|---|
| | Dim state | | Bright orange state | | Post-PA state with lactate(note 1) |
| | L-Lactate (+) | L-Lactate (−) | L-Lactate (+) | L-Lactate (−) | |
| Absorbance peak (nm) | 389 | 389 | 458 | 458 | 390 |
| Apparent EC(note 2) ($M^{-1}\cdot cm^{-1}$) | 2,800 | 3,200 | ND | 36,100 | 17,800 |
| Excitation peaks (nm) | 370/406 | 370/406 | 436/476 | 436/476 | 432/474 |
| Emission peak (nm) | 495 | 495 | 561 | 561 | 563 |
| Quantum yield of fluorescence | 0.15 | 0.088 | 0.48 | 0.48 | 0.41 |
| Relative Brightness | 0.013 | 0.0084 | ND | 0.52 | 0.22 |
| *In vitro* $\Delta F/F_{min}$ | + 0.27 | | + 0.047 | | NA |
| *In vitro* $\Delta F/F_{min}$ (PA) | NA | | NA | 5.8 | |
| *In vitro* $EC_{50}$ for $\Delta F/F_{min}$ (PA) (μM) | NA | | NA | 16.7 | |

**Table S1. Photophysical parameters of Paint-Lac1.** EC, extinction coefficient. NA, not applicable. ND, not determined. The two excitation peaks correlated with **Figure 3B** and the three-dimensional spectra in **Figure S16**. Quantum yields were determined under 460-nm excitation. Relative rightness is determined as apparent EC multiples quantum yield, then normalized by the brightness of EGFP

(**1**) The mixture of small population of photoactivated bright orange state and large population of non-photoactivated dim state.

(**2**) Apparent ECs were determined by $A_{458}$ / *c*, where $A_{458}$ is the absorbance at 458 nm and *c* is the concentration determined as previously reported[4]. Briefly, we assumed that the peak ECs of the NaOH-denatured dim-state Paint-Lac1 are equal to that of the NaOH-denatured GFP-type chromophore (44,000 $M^{-1}$ $cm^{-1}$). The *c* derived from the NaOH-denatured sample were used to determine all samples assuming the concentration in parallel experiments are strictly consistent.
